# Three-dimensional characterisation of osteocyte volumes at multiple scales, and its relationship with bone biology and genome evolution in ray-finned fishes

**DOI:** 10.1101/774778

**Authors:** Donald Davesne, Armin D. Schmitt, Vincent Fernandez, Roger B. J. Benson, Sophie Sanchez

## Abstract

Osteocytes, cells embedded within the bone mineral matrix, inform on key aspects of vertebrate biology. In particular, a relationship between volumes of the osteocytes and bone growth and/or genome size has been proposed for several tetrapod lineages. However, the variation in osteocyte volume across different scales is poorly characterised, and mostly relies on incomplete, two-dimensional information. In this study, we propose to characterise the variation of osteocyte volumes in ray-finned fishes (Actinopterygii), a clade including more than half of modern vertebrate species in which osteocyte biology is poorly known. We use X-ray synchrotron micro computed tomography (SRμCT) to achieve a three-dimensional visualisation of osteocytes and direct measurement of their volumes. Our specimen sample is designed to characterise osteocyte variation at three scales: within a bone, between the bones of one individual and between taxa spanning actinopterygian phylogeny. At the intra-bone scale, we find that osteocytes vary noticeably in volume between zones of organised and woven bone (being larger in the latter), and across cyclical bone deposition. This is probably explained by differences in bone deposition rate, with larger osteocytes contained in bone that deposits faster. Osteocyte volumes vary from one bone to another, for unclear reasons. Finally, we find that genome size is the best explanatory variable of osteocyte volume at the inter-specific scale: actinopterygian taxa with larger genomes (polyploid taxa in particular) have larger osteocytes. Our findings corroborate previous two-dimensional observations in tetrapods, and open new perspectives for actinopterygian bone evolution, physiology and palaeogenomics.

## Introduction

Osteocytes are the main cellular component of vertebrate bone and play important roles in bone growth and physiology (Hall, 2015). Through their ontogeny, the bone-producing cells called osteoblasts become embedded in the surrounding collagen matrix. From when they become enclosed within this mineralised tissue they are labelled as osteocytes, contained within interconnected voids called osteocyte lacunae. As such, lacunae provide a hard-tissue record of the osteocyte morphology (Franz-Odendaal *et al*., 2006; Bonewald, 2011; Webster *et al*., 2013), which varies according to that of the extracellular collagen matrix deposited during bone growth and remodelling (Amprino, 1947; Marotti, 1979; de Ricqlès *et al*., 1991; Kerschnitzki *et al*., 2011; van Oers *et al*., 2015). Therefore, the morphology of osteocyte lacunae has relevance to the study of bone development and osteocyte function among other topics. Furthermore, osteocyte lacunae are generally well-preserved in fossils, potentially enabling inference of various biological parameters that would otherwise be unknown in extinct animals, including developmental stage, metabolic and growth rates, the location of muscle insertion and genome size (Amprino, 1947; Francillon-Vieillot *et al*., 1990; de Ricqlès *et al*., 1991; D’Emic & Benson, 2013; Sanchez *et al*., 2013). However, the relationships between osteocyte morphology and many of these biological factors are not currently well constrained, so the potential of fossilised osteocyte lacunae to yield novel biological insights about extinct species is in doubt. This uncertainty is particularly relevant to deep patterns of physiological and genomic evolution in vertebrates, in particular on the ancestral lineages leading to speciose extant groups (e.g. birds, lissamphibians, teleost fishes; Organ *et al*., 2007, 2011; Laurin *et al*., 2016).

We present a multi-scale study that addresses this issue in actinopterygians (ray-finned fishes), a group that includes more than half of all vertebrate species today. Actinopterygian evolution saw key genomic events such as a whole-genome duplication specific to teleosts (Amores *et al*., 1998; Taylor *et al*., 2003; Hoegg *et al*., 2004; Donoghue & Purnell, 2005; Crow *et al*., 2006; Glasauer & Neuhauss, 2014) and several cases of polyploidy in smaller teleost subgroups (Leggatt & Iwama, 2003; Mable *et al*., 2011), as well as major physiological innovations such as the convergent evolution of various forms of endothermy (Block *et al*., 1993; Dickson & Graham, 2004; Wegner *et al*., 2015; Legendre & Davesne, in press). However, despite their importance, actinopterygians have received relatively little attention from a comparative histological perspective (Meunier, 2011).

Characterising the variation in osteocyte morphology is fundamental to constraining hypotheses about its biological causes, and can therefore provide insights into the multiple functions of osteocytes in living bone, currently not entirely understood in ray-finned fishes (Shahar & Dean, 2013; Doherty *et al*., 2015; Currey *et al*., 2017; Davesne *et al*., 2019). There are many accounts of variation in osteocyte morphology at different scales within various vertebrate lineages: (1) within a single bone (e.g. Canè *et al*., 1982; de Ricqlès *et al*., 1991; Cadena & Schweitzer, 2012; Sanchez *et al*., 2013), (2) among bones within an individual organism (e.g. D’Emic & Benson, 2013); (3) among individuals of different developmental stages within a species (e.g. (Sanchez *et al*., 2014); (4) among individuals of different species (inter-specific variation), even when considering homologous bones (e.g. D’Emic & Benson, 2013; Organ *et al*., 2016).

Variation in the volume of osteocytes is conspicuous, and is the focus of the present study. Various biological parameters have been proposed to correlate with osteocyte volume. In particular, bone growth rate has been widely discussed as a driver for this variation at the intra-bone and intra-individual scales (de Ricqlès *et al*., 1991; Remaggi *et al*., 1998; Cadena & Schweitzer, 2012; Stein & Prondvai, 2014). In contrast, other factors might be important at the inter-specific scale, including phylogeny, growth rate, basal metabolic rate, adult body size, and genome size (Organ *et al*., 2007; Montanari *et al*., 2011; D’Emic & Benson, 2013; Laurin *et al*., 2016). Which of these factors is most important can be determined by comparative statistical analysis of quantitative data on variation in osteocyte volume among species (e.g. in birds, D’Emic & Benson, 2013; Grunmeier & D’Emic, 2019). However, progress has been limited due to several challenges that affect most attempts to characterise osteocyte morphology in any detail. First, most existing studies are based on ground sections or thin sections, the traditional materials of bone histology. Depending on their thickness, these sections may preserve some information on the actual shapes and volumes of osteocyte lacunae. However, lacunae are more likely to be sectioned through than preserved complete, limiting the ability to characterise and quantify their threedimensional (3D) structure (Marotti, 1981; D’Emic & Benson, 2013; Stein & Prondvai, 2014). Furthermore, most studies so far have focused on a limited taxon diversity, most often humans (e.g., Marotti, 1979; Ardizzoni, 2001; Bromage *et al*., 2016) or model organisms (e.g., Remaggi *et al*., 1998; Schneider *et al*., 2007; Vatsa *et al*., 2008; Weigele & Franz-Odendaal, 2016; Suniaga *et al*., 2018; Ofer *et al*., 2019), and rarely use observations at multiple scales. Those that do include observations across several scales have used only small amounts of data (e.g., Cao *et al*., 2011; D’Emic & Benson, 2013). Finally, with a few exceptions (e.g. Kölliker, 1859; Stéphan, 1900; Amprino & Godina, 1956; Meunier, 1989; Meunier & Huysseune, 1992; Sanchez *et al*., 2014; Kamska *et al*., 2018; Davesne *et al*., 2019; Ofer *et al*., 2019), inter-specific comparisons of osteocyte morphology have been made only on tetrapods (four-limbed terrestrial vertebrates), representing only a subset of vertebrate diversity.

A small, but growing number of bone histological studies are based on high-resolution 3D imaging. Various methods have been used to three-dimensionally visualise osteocytes and/or their canalicular network, including confocal laser scanning microscopy (e.g., Vatsa *et al*., 2008; van Hove *et al*., 2009), serial focused ion beam and serial block-face scanning electron microscopy (e.g., Schneider *et al*., 2010), ptychographic X-ray computed tomography (Dierolf *et al*., 2010) and synchrotron X-ray micro- (SR-μCT) or nanotomography (Schneider *et al*., 2007; Pacureanu *et al*., 2012; Currey & Shahar, 2013; Mader *et al*., 2013; Sanchez *et al*., 2013, 2014, 2016; Dong *et al*., 2014; Peyrin *et al*., 2014; Suniaga *et al*., 2018; Kamska *et al*., 2018; O’Shea *et al*., 2019; Ofer *et al*., 2019), each method bringing comparative advantages and shortcomings (Sanchez *et al*., 2012; Webster *et al*., 2013; Goggin *et al*., 2016).

Our study aims to characterise and quantify the 3D structural variation of osteocyte lacunae, focusing on total size (volume), and using ray-finned fishes as a model system. This provides the first large-scale comparative study of variation in osteocyte volume among non-tetrapod vertebrates, facilitating future histological and palaeobiological inferences, and enabling broad inferences regarding osteocyte morphology and function in vertebrates.

## Material and methods

### Imaging method

Our dataset uses propagation phase contrast X-ray synchrotron micro computed tomography (PPC-SRμCT) to visualise osteocyte lacunae in 3D. This provides various advantages compared to other methods. Unlike thin sections, it allows direct three-dimensional visualisation of the shapes of lacunae and direct measurement of their volumes. The 3D datasets yielded by PPC-SRμCT also provide a convenient way to visualise and quantify lacuna abundance, orientation, and spatial distribution. Unlike thin sectioning and some other methods (e.g. electron microscopy), it is also entirely non-destructive. PPC-SRμCT also has many advantages over conventional X-ray μCT (Tafforeau *et al*., 2006; Sutton, 2008): (1) propagation phase contrast substantially increases the contrast between osteocyte lacunae and the surrounding bone matrix, and allows visualisation of structures that are otherwise invisible or difficult to trace (including osteocyte canaliculi and the cementing lines of reversal that delimit zones of remodelled secondary bone); (2) the parallel geometry and the power of the synchrotron beam allow imaging bone cell spaces in relatively large samples (which are several tens of centimetres) with a submicron voxel size; (3) due to the intensity of the synchrotron X-ray beam (i.e. brilliance), the signal-to-noise ratio is considerably improved and scan duration much shorter, reducing potential movement of the sample and therefore improving sharpness. Comparative studies have shown that PPC-SRμCT yields comparable results to physical thin sections (Sanchez *et al*., 2012) for visualisation of features such as osteocyte lacunae, lines of reversal, lines of arrested growth (LAGs) and vascularity. Shortcomings of any micron or sub-micron scale X-ray μCT based analysis include a relatively small field of view. The latter can be overcome by stitching (as commonly done with SEM, but here in 3D) at the expense of increasing the dataset size: with a common basic dataset size of ca. 15GB, doubling the scanned volume in all three dimensions results in a ca. 120GB dataset. Another drawback of the technique, compared to physical thin sectioning, is the increased difficulty to discriminate between bone types on the basis of the structure and orientation of collagen fibres, which are less visible than in physical sections. Measurements of a single lacuna can also be less precise with PPC-SRμCT than with physical sections or SEM, due to limitations of the resolution of tomograms and phase contrast artefacts that increase the margin of error (Sanchez *et al*., 2012). Nevertheless, the very high number of measurements that can be taken at once with this approach compensates for this. Crucially, PPC-SRμCT is by far the most cost- and time-effective method for 3D data acquisition (although data treatment is time consuming), especially when the study requires a large specimen sample to be analysed (Tafforeau *et al*., 2006).

### Specimen sample

We selected our specimen sample to address variation in osteocyte lacuna volume at the intra-bone, intra-skeletal and inter-specific scales. The specimens consist of dry bones of actinopterygians, sampled from skeletal preparations in museum collections (Table 1). We targeted two bones for most species due to their accessibility in prepared skeletons and unambiguous homology across ray-finned fishes: the left dentary (the main bone of the lower jaw) and an abdominal rib. While we collected dentaries for every species investigated, our rib sample covers only a subset of these species. The positional identities of the ribs were unknown in most cases, because we sampled them from disarticulated specimens. However, rib morphology is remarkably conserved across the abdominal region in most considered taxa, and we do not expect a large amount of histological variation to create a bias if non-homologous ribs are sampled for different taxa.

**Table 1:**
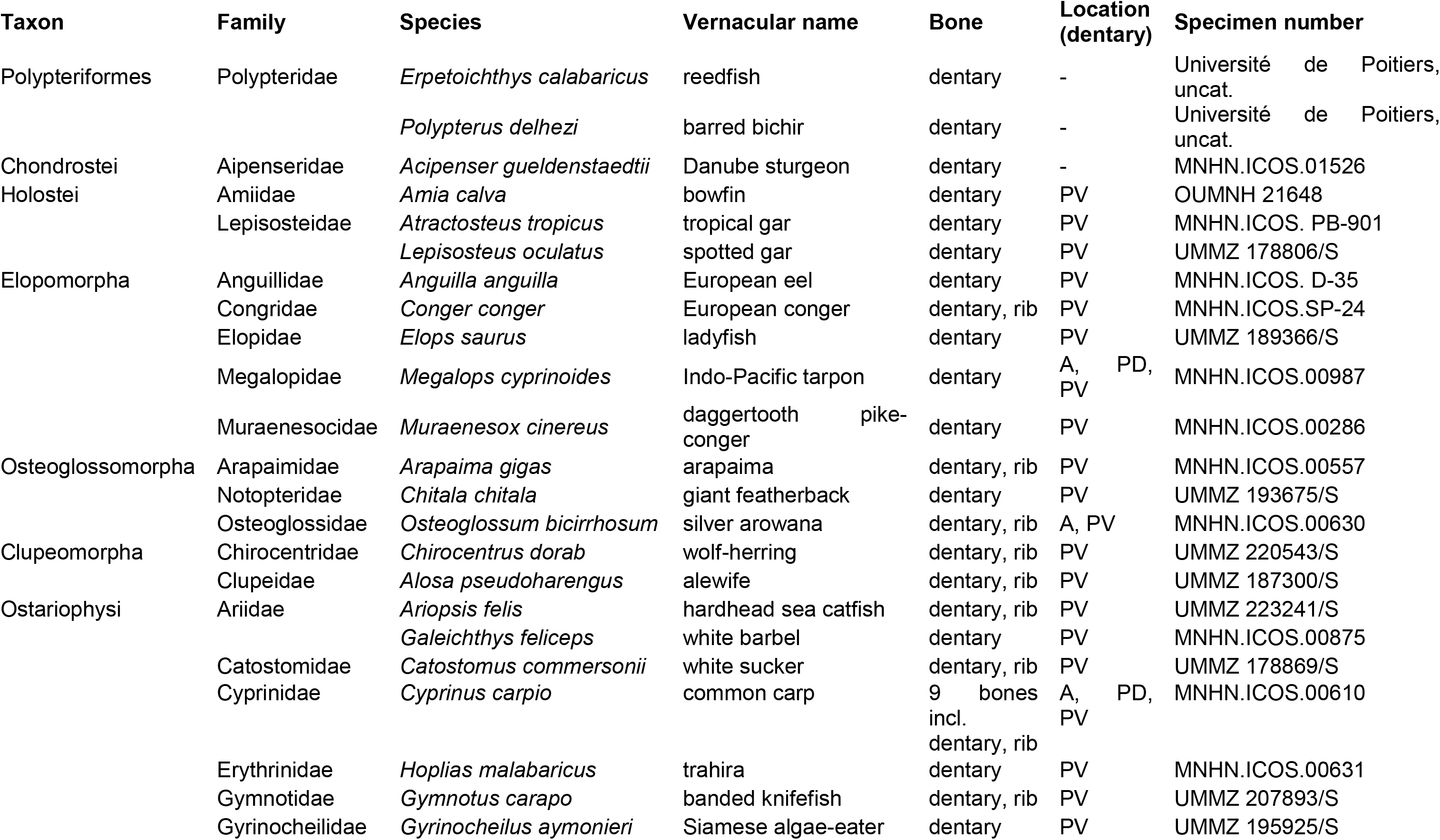
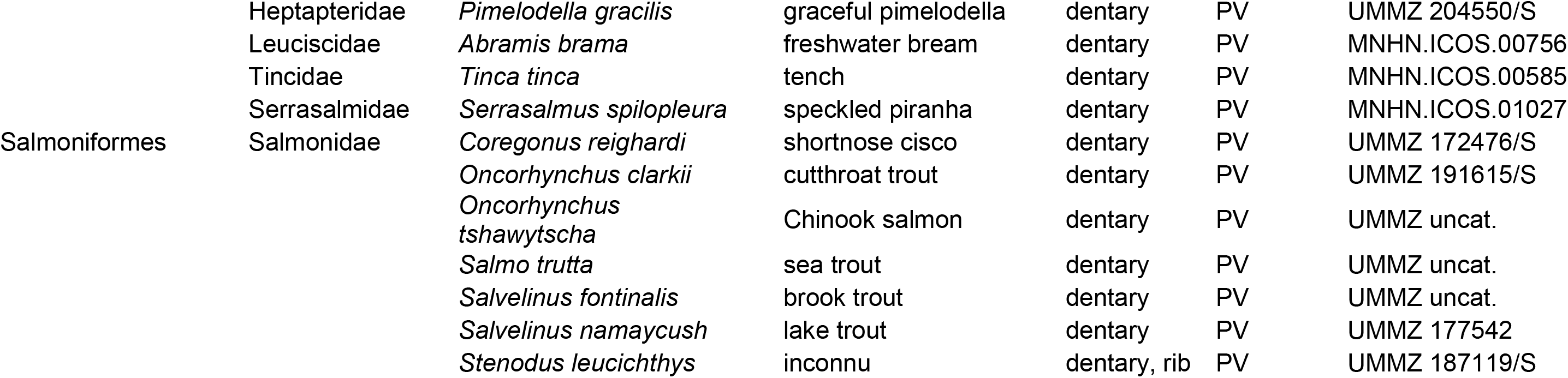
List of the specimens used in this study. The taxonomy follows Betancur-R. *et al*. (2017). Abbreviations for the location within the dentary: A, anterior portion of the dentary; PD, postero-dorsal branch; PV, postero-ventral branch. Institutional abbreviations: MNHN, Muséum national d’Histoire naturelle, Paris, France; OUMNH, Oxford University Museum of Natural History, Oxford, UK; UMMZ, University of Michigan Museum of Zoology, Ann Arbor, Michigan, USA.

To account for intra-bone variability, we acquired data on complete bones rather than using fragments. The selected specimens are large enough to guarantee that at least some amount of variation in bone tissue (e.g., woven vs. parallel-fibered bone; primary vs. secondary bone) is present due to the age of the individual, but small enough to allow for this variation to be incorporated into the scanned area. Larger specimens are also more likely to represent sexually mature individuals, although the exact age of the specimens is unknown in most cases.

To examine intra-skeletal variability, we characterised nine different bones from across the skeleton for one individual common carp (*Cyprinus carpio*). These bones represent several distinct functional and developmental units. In addition to a dentary (a dermal bone of the splanchnocranium) and an abdominal rib (an endochondral bone of the axial skeleton), we sampled a frontal bone (part of the neurocranium), an opercle, a maxilla, a pelvic bone, and a dorsal-fin spine (all dermal in origin), as well as the centrum of a caudal vertebra (of perichondral origin), and a pharyngeal bone (of mixed endochondral and dermal origin) corresponding to the fifth ceratobranchial (Eastman, 1971).

We adressed inter-specific variability by sampling taxa from the entire phylogenetic diversity of ray-finned fishes (Fig. 1). This includes non-teleost lineages Cladistia (bichirs), Chondrostei (sturgeons) and Holostei (gars and the bowfin), as well as teleost lineages Elopomorpha (eels, tarpons and relatives), Osteoglossomorpha (bony-tongues), Clupeomorpha (herrings and relatives), Ostariophysi (carps, characins, catfishes and relatives) and Salmoniformes (salmons and relatives). The clade Euteleostei includes approximately two-thirds of teleost diversity (Nelson *et al*., 2016), yet is only represented by salmoniforms in the sample. This is because euteleosts evolved ‘acellular’ (or anosteocytic) bone that is entirely devoid of osteocytes (Kölliker, 1859; Moss, 1961; Parenti, 1986; Meunier, 1989; Shahar & Dean, 2013; Davesne *et al*., 2018, 2019). Salmoniforms are one of the few exceptions, and may have re-acquired osteocytes secondarily during their early evolution (Davesne *et al*., 2019). The taxa we selected have known genome sizes (Gregory *et al*., 2007). Within each lineage, we chose taxa encompassing a broad range of adult body lengths and genome sizes (including polyploid taxa), to maximise the potential variability in osteocyte lacunar volumes.

**Figure 1:**
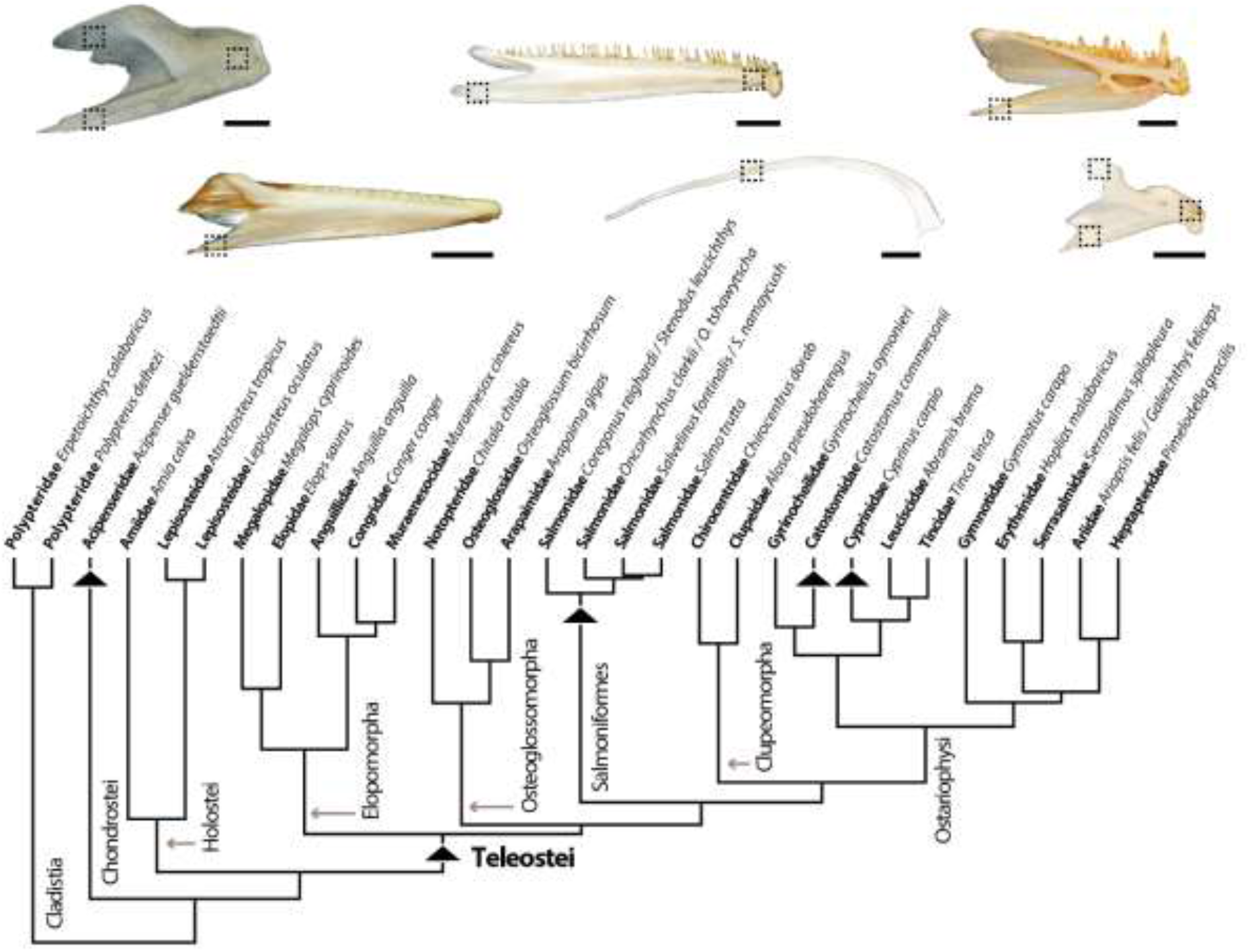
**(Top)** Examples of specimens used in the study. From left to right, top to bottom: left dentaries of the tarpon *Megalops cyprinoides* (MNHN.ICOS.00987), the arowana *Osteoglossum bicirrhosum* (MNHN.ICOS.00630), the trahira *Hoplias malabaricus* (MNHN.ICOS.00631), the conger eel *Conger conger* (MNHN.ICOS.PB-SP-24), abdominal rib and left dentary of the carp *Cyprinus carpio* (MNHN.ICOS.00610). Dashed lines symbolise the areas that we acquired with SRμCT. **(Bottom)** Time-calibrated phylogenetic tree of the species used in the study, obtained from the topology of Betancur-R *et al*. (2015). Several closely related species were represented by the same terminals in the analysis. Black arrowheads represent the lineages that underwent whole-genome duplications.

In total, our dataset includes 34 species (Table 1). 34 specimens were dentaries, ten were ribs, and for one species (*Cyprinus carpio*) nine different bones were collected.

### Data acquisition

All specimens were characterised at the ID19 beamline of the ESRF (European Synchrotron Radiation Facility, Grenoble, France) using PPC-SRμCT. Tomography acquisition was done with a setup producing volumes with an isotropic voxel size of 0.7 μm, adequate for measuring osteocyte lacunae. In order to accommodate to the different sizes and densities of our specimens, we used four different setups to acquire data (Supplementary Information), changing the energy (detected energies of 19, 35, 105 and 112 keV respectively) to adjust the transmission and the dose intake of the sample and to avoid movement during the scans (exposure to the beam could cause deformation of the sample in some of the setups we tried). Our setups used a pink beam from an undulator (at 19 and 35 keV) or filtered white beam from a wiggler (at 105 and 112 keV) and an indirect detector. Tomographic acquisitions consisted of 2999 projections with an exposure time ranging from 30 to 200 ms.

The volumes resulting from a single acquisition ranged from ca. 3 to 5 mm^3^, representing a small fraction of the entire bone. As we prioritised the taxonomic diversity of our sample rather than larger volume of a single bone, and because stitching μCT dataset can create unmanageable files sizes, we selected roughly homologous areas across our sample. We targeted a zone located in the postero-ventral branch of each dentary, and a zone located in the mid-shaft of each rib (Fig. 1). In addition, we targeted the anterior extremity of the dentary (just behind the symphysis) in three specimens (the carp *Cyprinus carpio*, the tarpon *Megalops cyprinoides* and the arowana *Osteoglossum bicirrhosum*), and the postero-dorsal branch in two specimens (*C. carpio* and *M. cyprinoides*) in order to evaluate variation between distinct areas within a single bone.

Tomographic reconstruction was performed using PyHST2 (Mirone *et al*., 2014), using the single distance phase retrieval approach (Paganin *et al*., 2002). The delta-beta value was set to 50 for all reconstructions, aiming to use the phase retrieval approach as a low pass filter and decrease the noise in data. The obtained 32-bits data were converted to 16-bits stack of jpeg2000 (compression of 10), using the 3D histogram from PyHST2 and a saturation of 0.002% of the 32-bits data. Additional processing included ring artefact correction (Lyckegaard *et al*., 2011).

Our PPC-SRμCT protocol yielded very high-quality images for most of our specimens, allowing to easily differentiate osteocyte lacunae from the surrounding bone matrix. In most cases, the scan resolution and phase contrast allow to distinguish very thin or faint structures such as osteocyte canaliculi and cementing lines of reversal, or in some cases, collagen lamellae (Fig. 2).

**Figure 2:**
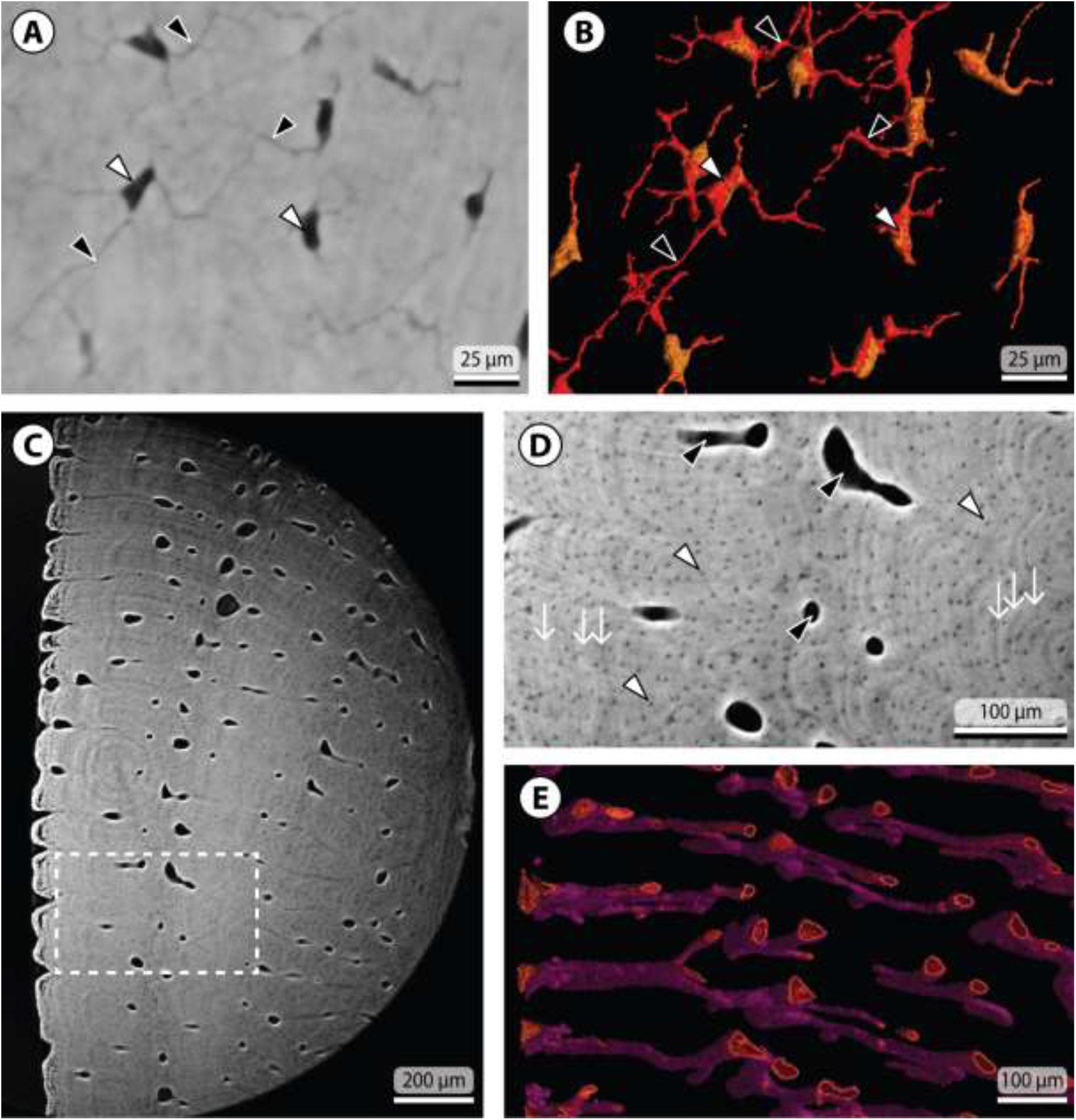
Examples of histological structures visible in our SRμCT tomograms. **(A)** Tomogram obtained from a dentary of the bichir *Polypterus delhesi* (Université de Poitiers, uncat.), clearly showing osteocyte lacunae (white arrowheads) and osteocyte canaliculi (black arrowheads). **(B)** Osteocyte lacunae (white arrowheads) and canaliculi (black arrowheads) segmented from the tomogram in (A). **(C)** Tomogram obtained from a dentary of the pike conger *Muraenesox cinereus* (MNHN.ICOS.00286), showing blood-vessel canals, osteocyte lacunae, and collagen lamellae. **(D)** Zoom on the inset in (C) showing osteocyte lacunae (white arrowheads), blood vessels (black arrowheads) and collagen lamellae (white arrows). **(E)** Blood vessels segmented from the tomogram in (C).

### Data visualisation and measurements

The tomograms (i.e. digital “slices”) obtained from our specimens were processed within VGStudioMAX v. 3.0 and 3.1 (Volume graphics, Heidelberg, Germany), to segment osteocyte lacunae from the surrounding bone matrix. We followed a standardised approach to facilitate inter-specimen comparisons and repeatability. The first step was to segment the internal and external bone surfaces, reflecting their location in the entire 3D volume. This was achieved by creating initial regions of interest (ROIs) encompassing volumes representing the internal cavities (e.g., blood-vessel canals) and the outside air. The region of interest was then inverted, yielding the shape of the bone as a new ROI. This inverted ROI was then duplicated twice, expanding one duplet by two voxels, and shrinking the other duplet by two voxels, using the “erode/dilate” function. Finally, the eroded ROI was subtracted from the dilated ROI, resulting in a hull volume with a wall thickness of 4 pixels, consisting of the dilated fraction minus the eroded fraction, i.e. the periosteal and endosteal bone surfaces. These segmented bone surfaces were useful to orient and interpret the internal histological data.

The second step was to segment osteocyte lacunae out of the bone matrix. We did not segment the canaliculi that accommodate cytoplasmic projections of the osteocytes, as they were not visible in every tomogram and thus could bias our estimates of the osteocyte lacuna volumes. Most of the lacunae are relatively homogeneous in terms of grey-value distribution, and were therefore straightforward to select and segment. We maintained consistent grey-value thresholds as far as it was possible, resulting in two ranges of thresholds. One was the strictest possible, keeping almost exclusively osteocyte lacunae but potentially underestimating their volume (“min threshold”); the second was a more lenient threshold (“max threshold”) that guaranteed that the whole lacuna was completely segmented, but potentially slightly overestimating their volumes and sometimes requiring additional manual postprocessing (due to other structures in close proximity that sometimes end up included within this threshold). In a given specimen, we always started by segmenting out osteocyte lacunae from the entire portion of bone that was imaged, enabling overall visualisation of variation in their volume, shape and orientation over a large portion of the bone. In most cases the imaged bone portion includes multiple types of bone and remodelled areas. We used bulk segmentations of osteocyte lacunae from the entire volume for visualisation purposes. However, these provide relatively imprecise and inconsistent measurements. One factor was the local tomography setup used (i.e. the imaged portion being significantly smaller than the surrounding bone), causing artificial variation in relative grey values, mostly manifested by a brightening towards the centre of the virtual slices (e.g. Fig. 2C). Furthermore, the presence of other small structures that were difficult to separate en-masse from osteocyte lacunae (e.g., canals of Williamson, muscle entheses, fibre bundles, post mortem cracks within the bone) also affected reliability of the measurements taken from the entire imaged bone portion. The measurements used in our analyses, therefore, were taken from more detailed segmentation of smaller, cuboid, regions of the image volume, which showed consistent within-region grey values allowing more consistent lacuna volume measurements. Detailed segmentation and manual editing of these smaller sample areas enabled to isolate accurately every included osteocyte lacuna. Lacunae that intersect with the edges of the cuboids were excluded.

Once osteocyte lacunae were segmented, we analysed them in the “Porosity/Inclusion” module of VGStudio MAX, to automatically measure the volume of each lacuna, as well as various other parameters (e.g., position in the xyz axis, length of the long axis). This was also used to provide visualisation, colouring lacunae according to their volume (with a consistent colour range across our sample) (Figs. 3–6).

**Figure 3:**
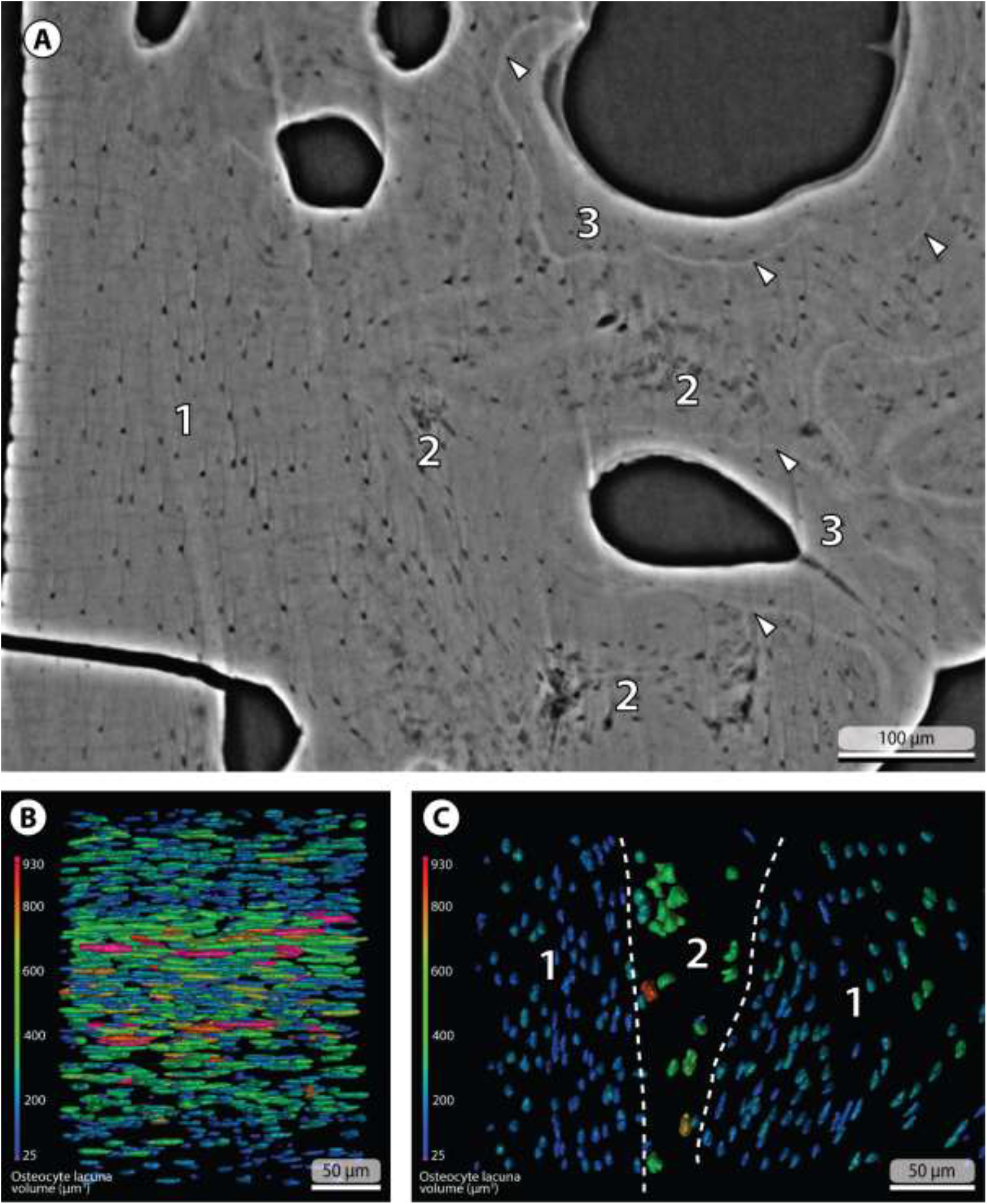
Examples of the different bone types we identify from our SRμCT tomograms and 3D reconstructions. **(A)** Tomogram obtained from a dentary (posteroventral branch) of the arowana *Osteoglossum bicirrhosum* (MNHN.ICOS.00630), showing the three main proposed bone types: organised bone with its osteocyte lacunae aligned and spaced regularly, mostly located in the outer portions of the bone (1); putative woven bone with a more irregular osteocyte lacuna pattern within a looser and heterogeneous extracellular matrix, located deep within the bone (2); secondary bone, delimited by cementing lines of reversal (white arrowheads), and located adjacent to blood vessels (3). **(B)** 3D reconstruction of the osteocyte lacunae from a region with an organised bone pattern in the pelvic bone of a carp *Cyprinus carpio* (MNHN.ICOS.00610). The lacunae are extremely organised, and all follow the same orientation. **(C)** 3D reconstruction of the osteocyte lacunae from the dentary (posteroventral branch) of a conger eel *Conger conger* (MNHN.ICOS.PB-SP-24). Lacunae in the inner area of putative woven bone (2) are larger, more irregular in shape and not aligned with each other, compared to the regular pattern of the lacunae in the areas of organised bone (1).

**Figure 4:**
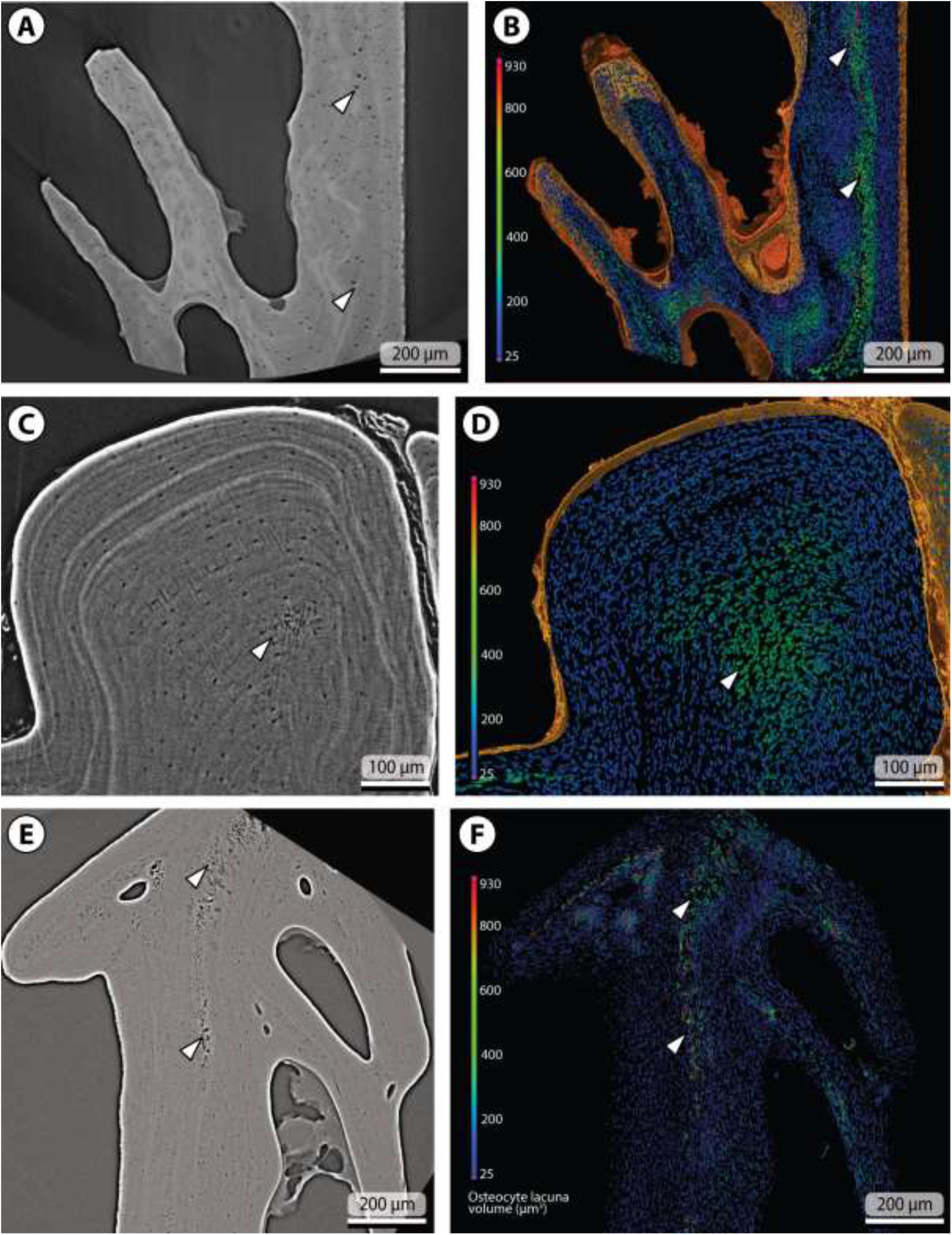
Examples of dentaries in which we infer the presence of woven bone (white arrowheads), characterised by larger osteocyte lacunae not organised in layers, and by a loose texture of the bone matrix. This woven bone is framed by an organised bone matrix that forms conspicuous layers of smaller osteocytes, resulting in a diploe structure of the bone tissue. **(A, B)** Dentary (posteroventral branch) of the conger eel *Conger conger* (MNHN.ICOS.PB-SP-24), detail of the tomogram and 3D reconstruction. **(C, D)** Dentary (posteroventral branch) of the ladyfish *Elops saurus* (UMMZ 189366/S), detail of the tomogram and 3D reconstruction. **(E, F)** Dentary (posteroventral branch) of the trahira *Hoplias malabaricus* (MHNH.ICOS.00631), tomogram and 3D reconstruction, presumably showing two folds of the bone fused to each other (woven bone is found at the junction of these two folds).

**Figure 5:**
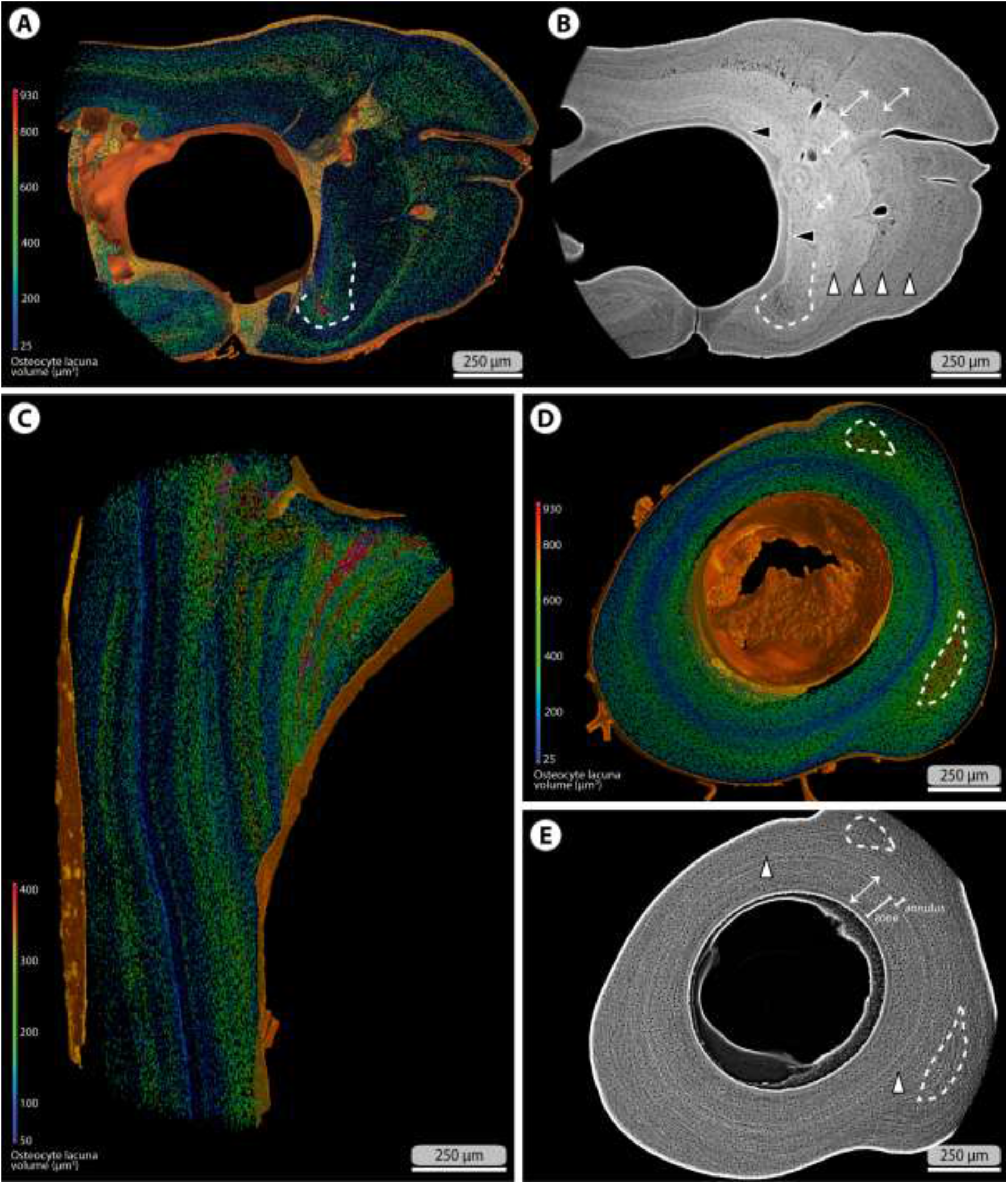
Examples of specimens in which alternating layers of osteocyte lacunae of varying volumes are well visible within the periosteal bone deposit. **(A)** 3D reconstruction of a rib from a carp *Cyprinus carpio* (MNHN.ICOS.00610) and **(B)** the associated tomogram. A putative area of woven bone (white dashed line) is characterised by larger osteocyte lacunae, and a looser bone matrix. A cementing line of reversal (black arrowheads), delimitating a layer of secondary bone covering the endosteal surface, is visible. At least four growth marks (white double arrows) are visible, each including a zone of relatively fast growing bone and an annulus of slower growing bone. Each growth mark is followed by a line of arrested growth (white arrowheads). The zones and annuli are materialised by larger and smaller osteocyte lacunae, respectively. In addition, osteocyte volume varies from one growth mark to another. **(C)** 3D reconstruction of the operculum from an arowana *Osteoglossum bicirrhosum* (MNHN.ICOS.00630). Note the different colour scale for the osteocyte volumes, exacerbating volume alternation. Whether this variation reflects growth marks or not is unclear. **(D)** 3D reconstruction of a rib from an arowana *Osteoglossum bicirrhosum* (MNHN.ICOS.00630) and **(E)** the associated tomogram. One growth mark is visible (white double arrow), that includes a well-visible concentric layer of smaller osteocytes that we interpret as an annulus predating a line of arrested growth (white arrowhead). Two putative areas of woven bone (white dashed line) are locally characterised by larger osteocyte lacunae, and probably associated with an increase in growth rate related to rib morphogenesis.

**Figure 6:**
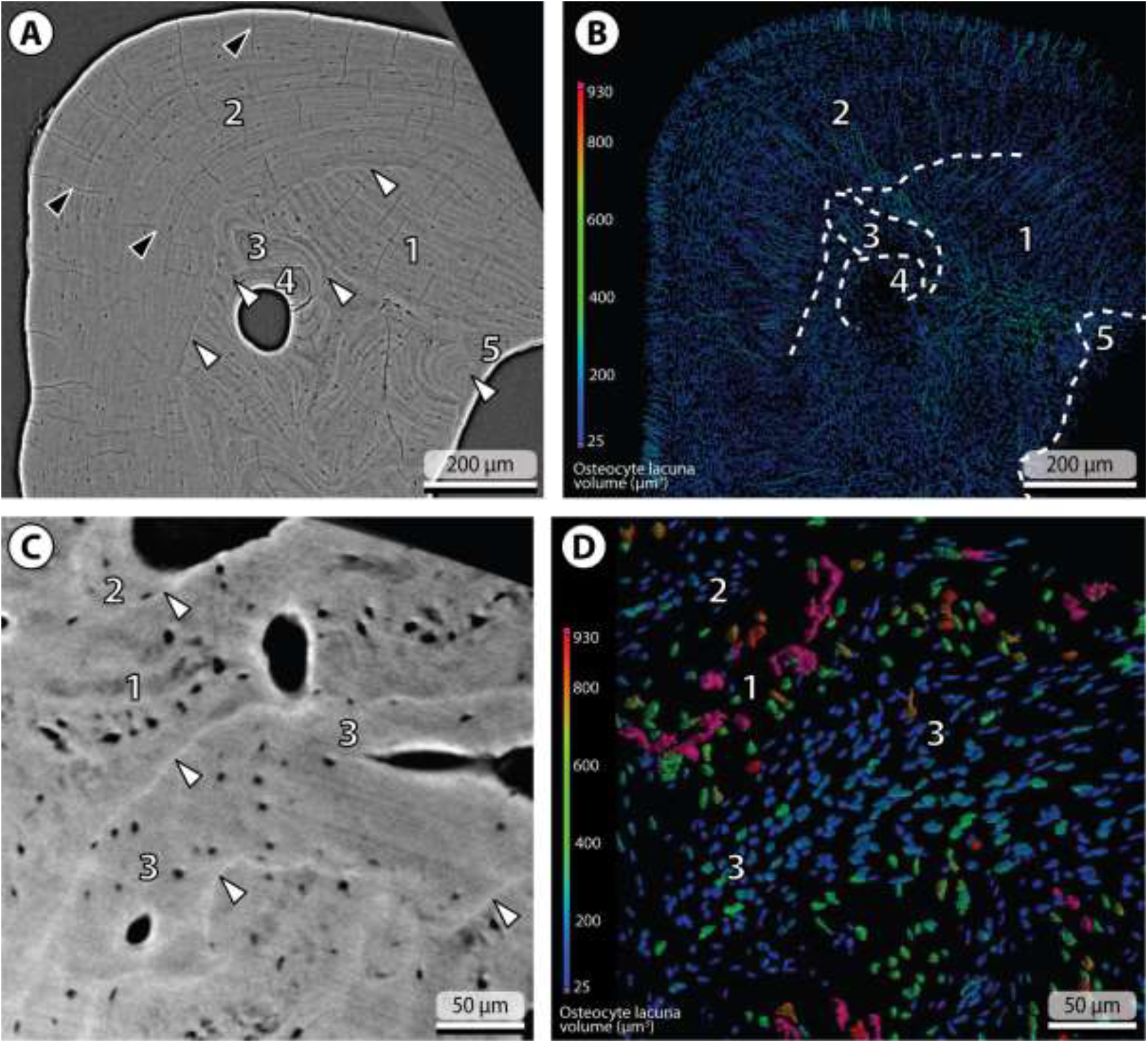
Examples of specimens in which cementing lines of reversal (white arrowheads in A, C and dashed lines in B) delimitating areas of secondary bone are visible. **(A)** Tomogram of a dentary (posteroventral branch) from a gar *Lepisosteus oculatus* (UMMZ 178806/S) and **(B)** the segmented osteocyte lacunae and canals of Williamson. The cementing lines document successive steps of bone growth: from a zone of primary, potentially lamellar bone (1), the resorption line delimitates more recent bone, growing in a different orientation (2). Further resorption yields successive waves of secondary (3 and 4) bone growth close to a blood-vessel canal. More secondary bone is visible on the lingual surface of the dentary (5). The tomogram highlights the lamellar organisation of all these successive bone deposits, thereby suggesting that all of them are organised tissue types (probably of lamellar nature). The volumes of osteocyte lacunae do not vary significantly between the areas of primary and secondary bone. Also note the abundant canals of Williamson (black arrowheads in A), most of them having been segmented alongside osteocyte lacunae (B). **(C)** Tomogram of a dentary (anterior extremity) from a carp *Cyprinus carpio* (MNHN.ICOS.00610) showing a region of putative woven bone (1) and deposits of secondary organised bone (2 and 3) and **(D)** the segmented osteocyte lacunae. Cementing reversal lines separate the primary bone (1) from at least two areas of secondary bone (2, 3). In this specimen, the osteocyte lacunae of secondary organised bone appear to be much smaller than in the primary woven bone (D).

### Bone pattern identification

From this visualisation of osteocyte lacuna volumes, considered alongside the corresponding tomograms, we detect consistent patterns of variation across our specimens. Two main patterns are identified, as follows:

(1) In both our tomograms (Fig. 3A) and their 3D reconstructions (Fig. 3B), we observe successive layers of well-organised osteocytes that are evenly spaced, all aligned with each other and with an elongate morphology. This pattern (‘organised bone’) is mostly found close to the outer (i.e. periosteal and endosteal) bone surfaces. In tomograms, it is also associated to a homogeneous bone matrix, sometimes with visible collagen lamellae (e.g. Figs. 2C-D, 4C, 4E, 6A).

(2) In areas located deep within the bone, a second pattern (‘woven bone’, as defined below) is characterised by a greater variability in osteocyte orientations, shapes, and spatial distribution (Fig. 3C). Osteocytes that are more globular than elongate in shape are mostly found in these areas (Fig. 3C). In addition, these areas show different colour and texture in tomograms, specifically darker grey values probably denoting a lower density and degree of organisation of the matrix (Fig. 3A). Classical bone typology distinguishes several bone matrix categories, namely woven, parallel-fibered and lamellar bone (Amprino, 1947; Francillon-Vieillot *et al*., 1990; de Ricqlès *et al*., 1991; De Margerie *et al*., 2002). These categories are determined by variations in growth rates, where woven bone presents the lowest degree of organisation in its extracellular matrix, indicative of a faster growth, while parallel-fibered and lamellar bone are deposited more slowly and show the most regular and organised extracellular matrix (with the collagen fibres organised parallel or perpendicular to each other). Woven bone develops early and is then located further away from the bone surfaces, or locally at junctions between two folds of a bone (de Ricqlès *et al*., 1991). Parallel-fibered and lamellar bones deposit later and are then found under the bone surfaces (periosteal and endosteal). In more recent studies, a distinction has been made between static and dynamic osteogenesis (Ferretti *et al*., 2002), with woven bone derived from static osteogenesis, while parallel-fibered and lamellar bone derive from dynamic osteogenesis (Prondvai *et al*., 2014). While collagen fibres are not typically visible with μCT data, a few of our tomograms do appear to show these (e.g., Fig. 2C, D). Osteocyte elongation follows the orientation of the extracellular collagen fibres (Kerschnitzki *et al*., 2011), meaning that the osteocytes of parallel-fibered and lamellar bone are oriented parallel to each other and organised in regular layers. However, there is no distinction between parallel-fibered and lamellar bone on the basis of osteocyte morphology alone (e.g. Warshaw *et al*., 2017). For the sake of identifying bone types from patterns of osteocyte morphology and distribution, it is then preferable to lump these two categories together as “organised bone” (Stein & Prondvai, 2014), since even though parallel-fibered bone seems to be the dominant organised bone type in actinopterygians (de Ricqlès *et al*., 1991), lamellar bone can also be found (e.g. Fig. 2C-D). We then propose that there is a correspondence between bone matrix categories and the two patterns of osteocyte morphology and distribution we identified: they represent organised bone (either parallel-fibered or lamellar) and woven bone (Figs. 3, 4).

In addition, we identify areas of bone, most often located adjacent to bloodvessel canals, and delimited by cementing lines of reversal. The latter are visible in SRμCT tomograms thanks to their conspicuous brightness (Figs. 3A, 6). This is due to their higher apparent density, most likely related to their relatively lower collagen and higher mineral content (Schaffler *et al*., 1987; Skedros *et al*., 2005). We interpret these cementing lines of reversal as delimitating secondary bone, deposited during bone modelling or remodelling. Secondary bone can be either woven or organised, depending on its rate of deposition. However, osteocytes we found in secondary bone correspond more to our organised bone pattern (Figs. 3A, 6), suggesting that secondary bone in the taxa we studied is either parallel-fibered or lamellar and deposits slowly, rather than woven and quickly deposited.

## Comparative observations and analyses

### Intra-bone variation

Two bone patterns corresponding to organised and woven bone were recognised on the basis of their cellular organisation (see above, ‘Bone pattern identification’). Here we characterise the volumes of osteocytes within bone of these patterns in the skeleton of actinopterygians. Several studies suggested differences in cell volumes for different bone types (e.g., Remaggi *et al*., 1998; Cadena & Schweitzer, 2012; Sanchez *et al*., 2013; Grunmeier & D’Emic, 2019). However these differences have so far been almost exclusively characterised in sarcopterygians (and most of the time, in mammals). We highlight the differences in 3D osteocyte lacuna volume in actinopterygians, between primary organised bone, primary woven bone, and secondary organised bone by qualitative comparison of measured cell volumes.

We also compared the volumes of osteocyte lacunae in the compact, primary bone of dentaries and ribs. Since we scanned a zone located in the postero-ventral process of dentaries, bone was relatively thin, often allowing to encompass its whole thickness. This facilitated the interpretation of the histological structures we observed (Figs. 3–6). In ribs, we considered the entire thickness of the compact bone surrounding the central cavity (Fig. 5).

Finally, we also asked whether different regions of the same bone showed differences in osteocyte lacuna volume independent of these other factors by scanning multiple separate regions of the dentary (anterior extremity, postero-dorsal process, and postero-ventral process) in specimens of the carp *Cyprinus carpio*, the tarpon *Megalops cyprinoides* and the arowana *Osteoglossum bicirrhosum*.

### Intra-skeletal variation

To evaluate variation in osteocyte lacuna volumes within the skeleton of a single individual, we segmented osteocyte lacunae from nine different bones of a carp *Cyprinus carpio* (see above, ‘Specimen sample’), and from the rib and dentary of nine other actinopterygian species. We took a cuboid subsample from each of these bones, targeting primary compact bone. This allowed comparison of osteocyte lacuna volumes between different skeletal elements, which we assessed by visual appraisal of the median and interquartile range. This allowed us to place into context our observations of variation in osteocyte lacuna volume among bone types and between different regions of the dentary into context and ask which of these three sources of variation in osteocyte volume has the largest effect size.

### Inter-specific variation

At the inter-specific level, osteocyte lacuna volumes have been proposed to correlate with various biological parameters. D’Emic & Benson (2013) and Grunmeier & D’Emic (2019) tested this scaling for modern birds in bone derived from both static and dynamic osteogenesis, using various biological parameters including body mass, growth rate, mass-specific metabolic rate, and genome size. They found that growth rate and, most notably, genome size had a statistically significant relationship with osteocyte lacuna volume.

For our actinopterygian dataset, we evaluated the relationships between osteocyte lacuna volumes and a series of explanatory variables using phylogenetic generalised least squares regression (pGLS; Grafen, 1989). These analyses were implemented in R version 3.6.0 (R Core Team, 2019) using functions from the packages nlme version 3.1-139 (Pinheiro *et al*., 2019) and ape version 5.0 (Paradis & Schliep, 2019). We allowed λ, a measure of phylogenetic signal in the relationship between variables (Pagel, 1999), to vary, and its value was estimated during fitting of the regression models. Estimated values of λ close to zero indicate the absence of phylogenetic signal, in which case the analysis replicates ordinary least squares regression. Values of λ close to 1.0 indicate strong phylogenetic signal and replicate pGLS assuming Brownian motion (Felsenstein, 1985). This analysis makes use of a phylogeny with branch lengths. For this, we used the time-tree of Betancur-R. *et al*. (2015).

Osteocyte volumes span a range of values within each region of interest that we measured. We used the median values of these measurements in each of our analyses, and performed separate analyses for the “min” and “max” segmentation threshold values (see above, Material and methods – Data treatment and measurements). The measurements for median osteocyte volumes are available online in the Dryad Digital Repository (URL upon acceptance). We fit models explaining log_10_-transformed osteocyte lacuna volumes with the following explanatory variables: species genome size, body length and ploidy level, plus every combination of these three variables. Akaike’s information criterion for finite sample sizes (AICc; Sugiura, 1978) was used to compute Akaike weights (Burnham & Anderson, 2002). R^2^ values for models were computed by comparison to an intercept-only null model using the generalised formulation of Nagelkerke (1991). Maximum body lengths of sampled species were collected from FishBase (Froese & Pauly, 2018), and may act as a proxy of growth rate (we were not able to access reliable direct measures of growth rate). We used this metric because other size measures were only available for a subset of our sampled species (maximum body mass, common body length and mature body length), but comparisons with other measures are available (Figs. S1-4) and showed similar results. We obtained genome sizes from the Animal Genome Size Database (Gregory, 2019). Multiple measured genome sizes are available for some species, and for these we used the median value. For two taxa, the gar *Atractosteus tropicus* and the bichir *Polypterus delhesi* information on genome size was not available, so we used that of closely related members of the same genera *A. spatula* and *P. palmas*. Estimates of metabolic rate are rare for actinopterygians, with a few exceptions (e.g., Clarke & Johnston, 1999), and we did not find any for the vast majority of our taxon sample, thus excluding this parameter from the analyses. The original values of body length, body mass and genome size we used are available online in the Dryad Digital Repository (URL upon acceptance).

Comparative analyses were run separately for the osteocyte lacunae measured from the dentaries and from the ribs. The species that we sampled for our study do not map precisely to the tips of the phylogeny used for our comparative analyses (Betancur-R. *et al*. 2015). For our sample of dentary osteocyte volumes, complete data were available for 34 species, which mapped to 30 unique terminals (Fig. 1) in the phylogeny of Betancur-R. *et al*. (2015). In several cases, multiple species in our dataset corresponded to a single tip in the phylogeny (e.g. either *Arius felis* or *Galeichthys feliceps* could correspond to the marker taxonomic unit for Ariidae; Fig. 1). Therefore, we performed 100 iterations of each analysis, drawing taxa to tips of the phylogeny at with equal probabilities per iteration. This resulted in a sample size of N = 30 for each iteration of our analysis for dentary osteocyte lacuna volumes. For our sample of rib osteocyte volumes, complete data were available for N = 10 species, all of which mapped to unique tips in the phylogeny of Betancur-R. *et al*. (2015).

## Results

### Intra-bone variation

We find conspicuous variation in osteocyte lacuna volumes, consistently across our sample. Each bone shows substantial, spatially structured variation in osteocyte lacuna volumes, and this variation seems to be partitioned among bone patterns described above (‘Bone pattern identification’), with differences between primary organised bone, primary woven bone, and secondary organised bone.

(1) Osteocyte lacunae in regions of woven bone are generally larger than in the adjacent organised bone (Figs. 3C, 4). This volume difference can be important, ranging from four-fold (~100 μm^3^ in organised bone, ~400 μm^3^ in woven bone in the ladyfish *Elops saurus;* Fig. 4D) to six-fold or more (less than 100 μm^3^ in organised bone, more than 600 μm^3^ in woven bone in the trahira *Hoplias malabaricus*; Fig. 4F).

(2) Successive layers of larger and smaller osteocytes are found in the organised bone of most ribs (Fig. 5A, D) and dermal elements, including dentaries (Fig. 5C). The number of apparent layers varies among specimens, ranging from one main layer of smaller osteocytes separating two layers of larger ones (Fig. 5D) to multiple successive layers of varying osteocyte lacuna volumes (Fig. 5A, C). It is likely that this variation reflects different types of variation in growth rates through time, possibly including seasonal or annual cycles.

(3) There is no consistent pattern of variation in osteocyte lacuna volumes between primary and secondary bone. In some cases, osteocyte lacunae are approximately of the same median size between these bone types (Fig. 6B). In others the difference is striking (Fig. 6D), in which cases osteocytes tend to be smaller in secondary bone compared to primary bone (Fig. 6D).

### Intra-skeletal variation

Osteocyte lacuna volumes in the carp *Cyprinus carpio* vary among elements of the skeleton (Fig. 7). Most elements have median osteocyte lacuna volumes ranging between 167 μm^3^ (in the maxilla) and 253 μm^3^ (in the frontal), with strongly overlapping distributions (Fig. 7). The dorsal-fin spine (507 μm^3^) shows much larger osteocytes, such that osteocyte sizes span approximately 3.5-fold range among bones of the skeleton. Variation in osteocyte volume does not seem to reflect the ontogenetic origin of the bones: relatively larger osteocytes are found in both endochondral bones such as the rib and in dermal bones such as the dorsal-fin spine. There is also no obvious correlation with the sequence of ossification: bones that ossify earlier in the zebrafish *Danio rerio* – a cypriniform closely related to *C. carpio* – such as the opercle and vertebrae (Cubbage & Mabee, 1996), do not have notably larger or smaller osteocytes than the other bones.

**Figure 7:**
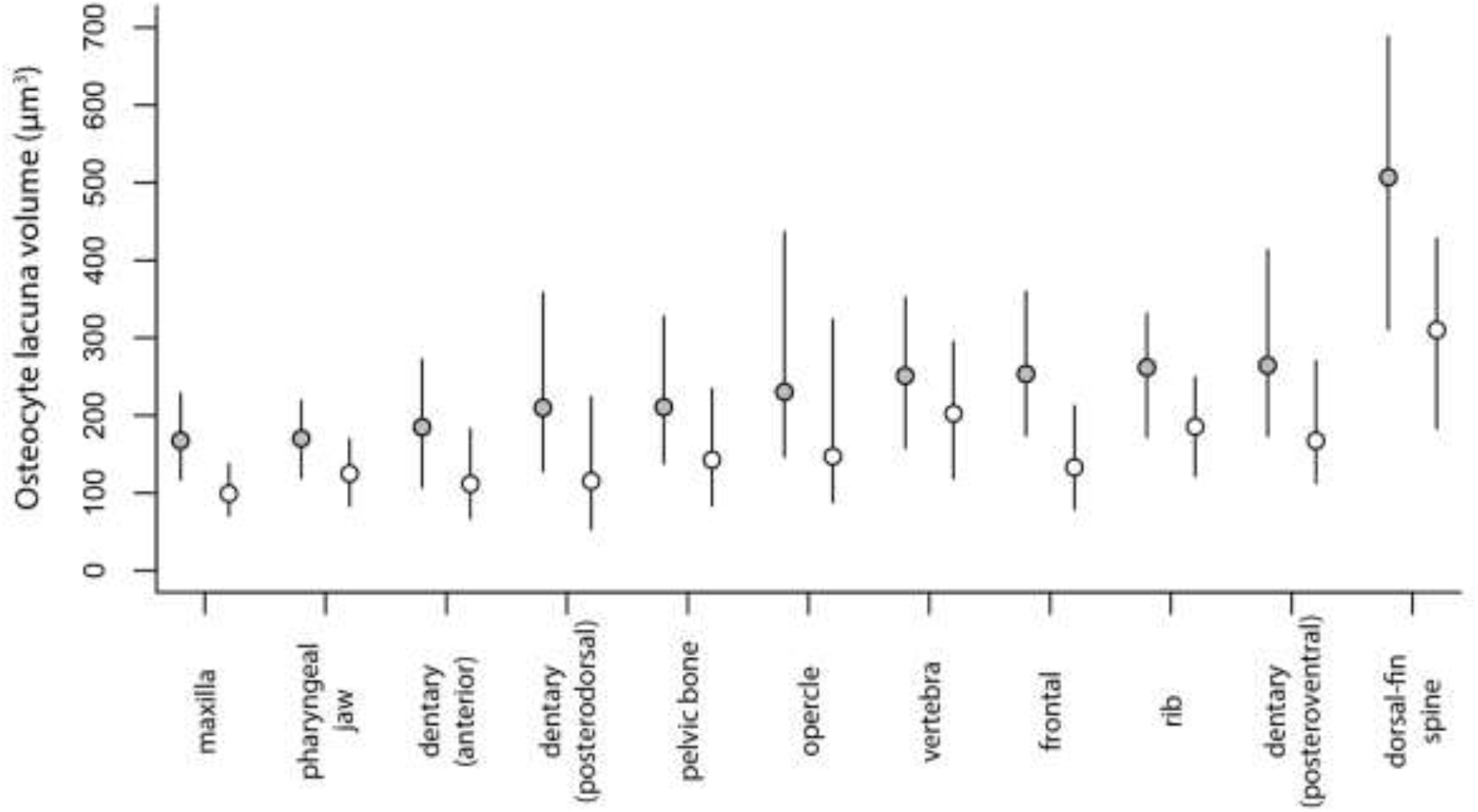
Comparison of the osteocyte lacuna volumes between nine different bones of a single carp (*Cyprinus carpio*) individual, with the medians and interquartile ranges shown. “Min threshold” in white, “max threshold” in grey.

Compared to the variation observed between different bones of the same skeleton, lacunae from different areas of the same bone yield highly overlapping and similar measurements within an individual dentary: postero-dorsal and postero-ventral processes, and anterior extremity behind the symphysis in *Cyprinus carpio, Megalops cyprinoides* and *Osteoglossum bicirrhosum* (Fig. 8).

**Figure 8:**
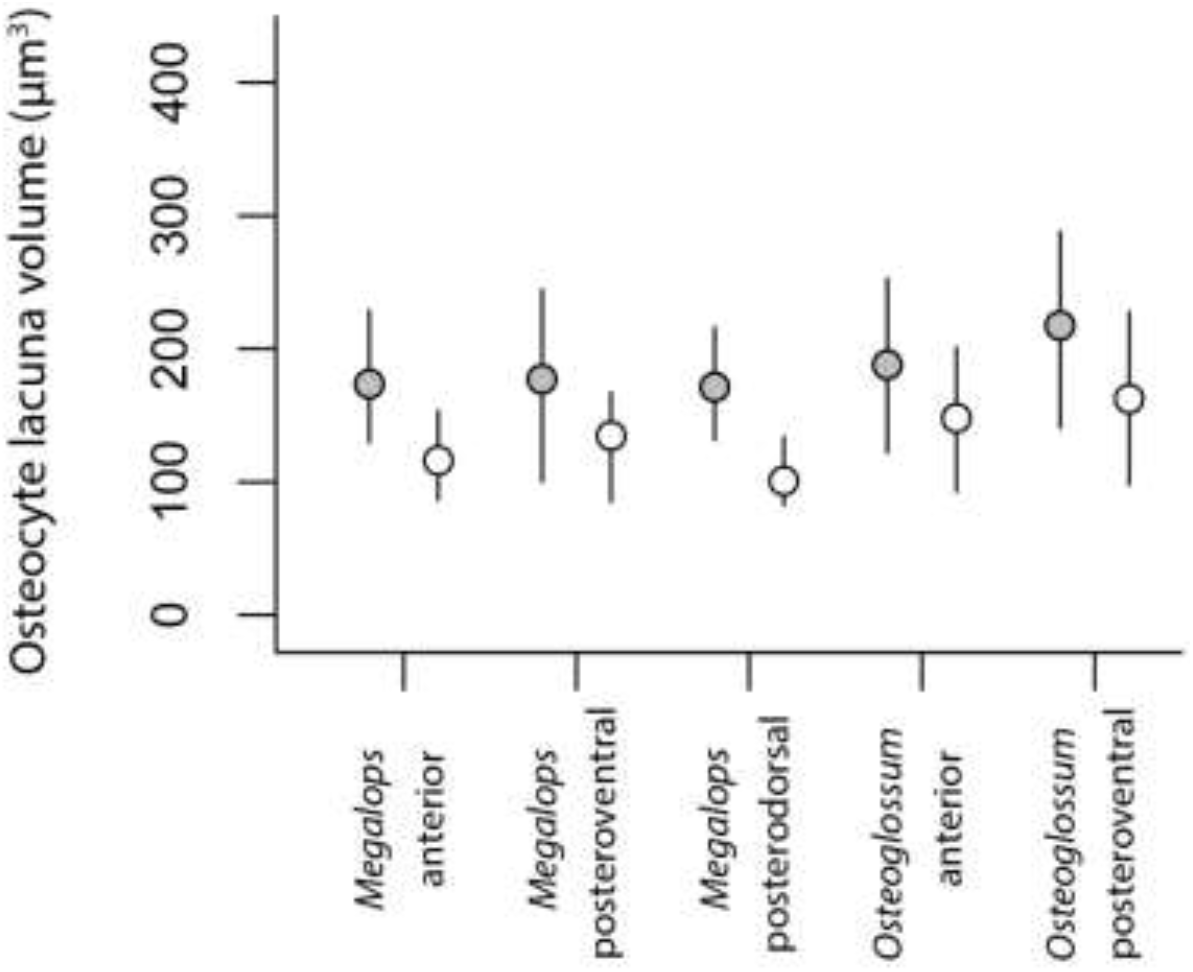
Comparison of the osteocyte lacuna volumes between different areas of the dentary in a tarpon (*Megalops cyprinoides*) and an arowana (*Osteoglossum bicirrhosum*), with the medians and interquartile ranges shown. For the carp (*Cyprinus carpio*), see Fig. 7. “Min threshold” in white, “max threshold” in grey.

Finally, we observe some variation in the relative sizes of osteocytes in dentaries compared to ribs in the same individual across the ten species for which these data are available. Many species show highly similar osteocyte volumes in primary lamellar bone of the dentary and rib (Fig. 9). However, the salmoniform *Stenodus leucichthys* and the carp *C. carpio* show a substantial difference between the sizes of osteocytes in these elements within an individual. Osteocytes of the dentary of *S. leucichthys* are much larger than those of the rib, whereas osteocytes of the dentary are smaller in *C. carpio* (Fig. 9).

**Figure 9:**
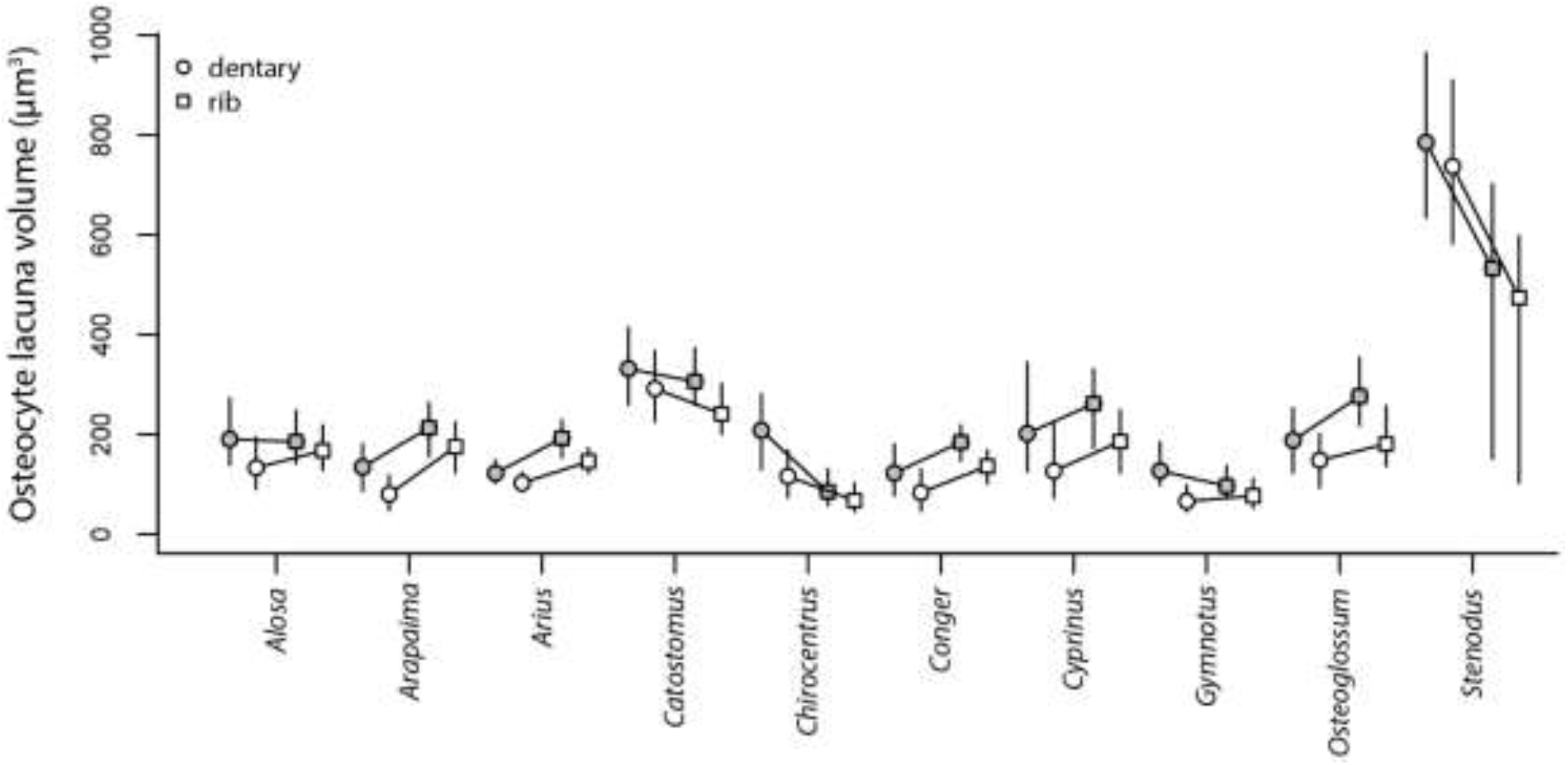
Comparison of the osteocyte lacuna volumes between ribs (squares) and dentaries (circles) in ten teleost species, with the medians and interquartile ranges shown. “Min threshold” in white, “max threshold” in grey.

### Inter-specific variation

Our phylogenetic regression analyses indicate that osteocyte lacuna volumes correlate with genome sizes and ploidy levels, both for ribs and dentaries (Fig. 10). Information criteria (AICc) provide equally strong support for models explaining median osteocyte lacuna volume using either genome C-size or ploidy level, and reject models that include multiple explanatory variables, and those including maximum species body length (Tables 2, S1).

**Figure 10:**
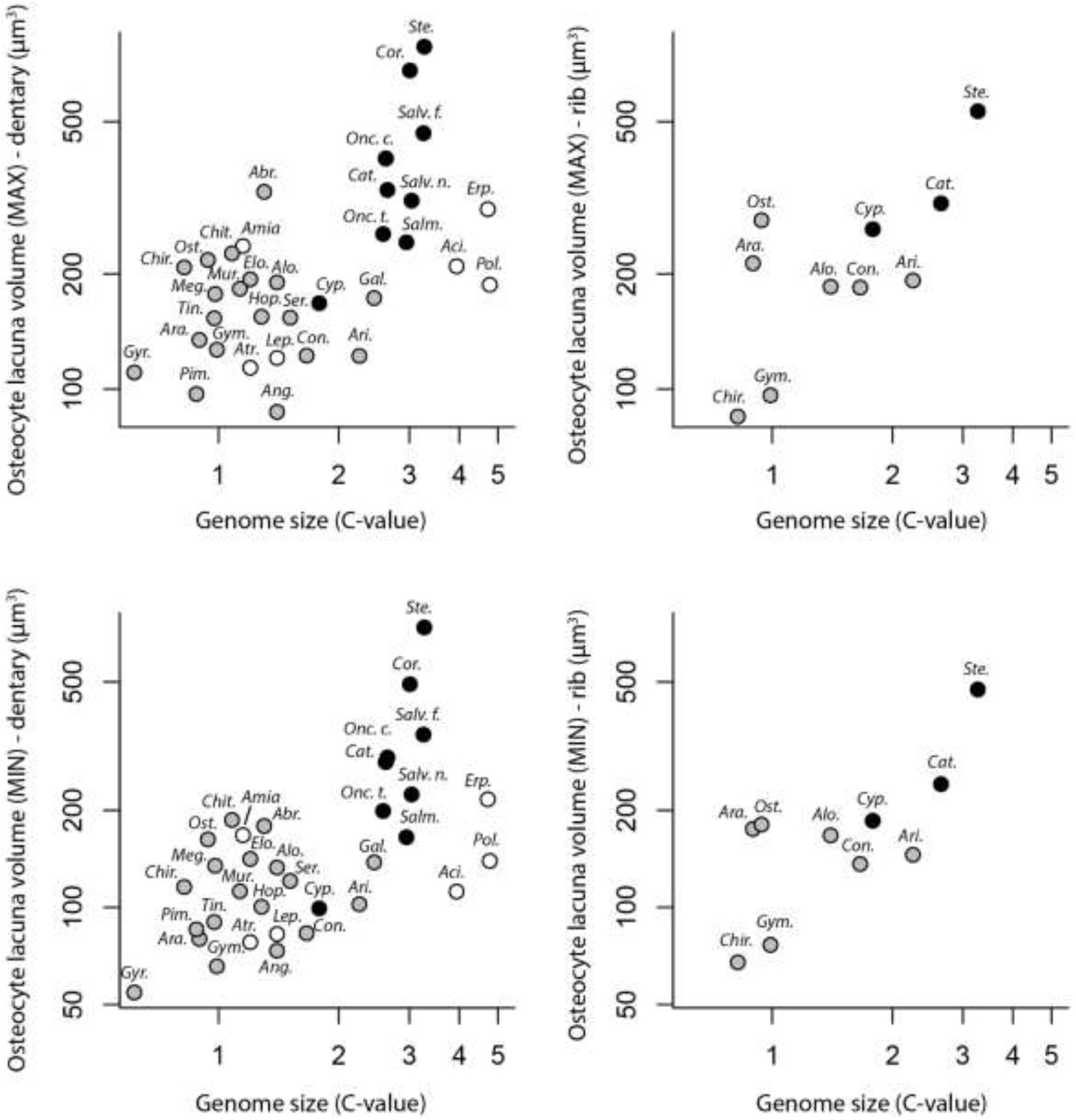
Phylogenetic generalised least squares regression (pGLS) of median osteocyte lacuna volumes against genome sizes (both log_10_-transformed). Analyses were ran separately with the dentary (left column) and the rib (right column) datasets on one hand, and using the “max threshold” (top row) and “min threshold” (bottom row) on the other hand. Non-teleost taxa are in white, recent polyploid teleosts are in black, non-polyploid teleosts taxa are in grey.

**Table 2:**
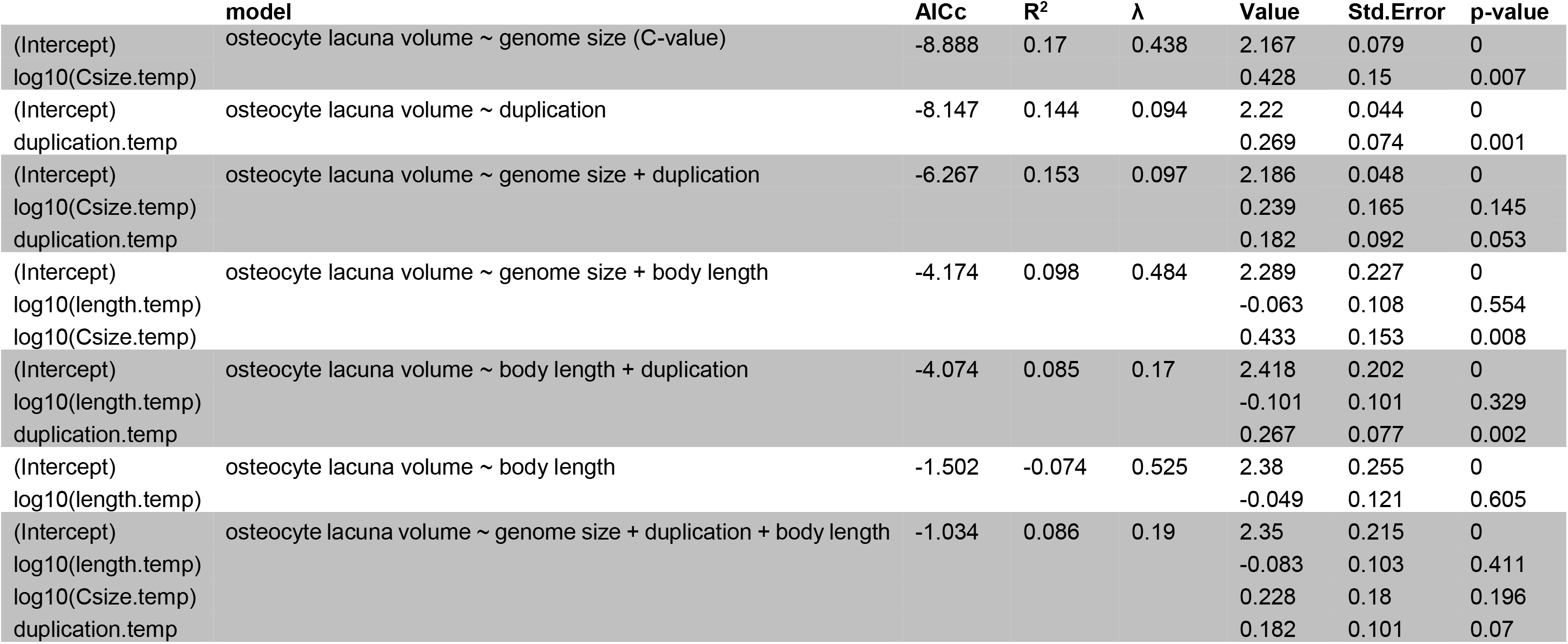
Results of the phylogenetic generalised least squares regression (pGLS). 100 iterations have been performed, and the results presented here are for the 50% quantile of the analysis on dentaries and at the “max” threshold. For the other series of results, see Table S2.

Indeed, maximum species body length does not correlate with osteocyte lacuna volume when analysed alone (Tables 2, S1). Although genome size and ploidy levels are individually correlated with osteocyte lacuna volumes, these variables have nonsignificant coefficients in models that include both together as explanatory variables, indicating that these variables contain redundant information.

The correlation with genome size has an R^2^ value of 0.17 (dentary, “max” threshold for osteocyte lacuna segmentation and measurement), 0.24 (dentary, “min” threshold), 0.61 (rib, “max” threshold), or 0.50 (rib, “min” threshold). The correlation with ploidy level has an R^2^ value of 0.14 (dentary, “max” and “min” thresholds), 0.30 (rib, “max” threshold), or 0.05 (rib, “min” threshold, a poorly-supported model according to AICc; Table S1). Phylogenetic signal in these relationships is intermediate, ranging from λ = 0.36 to λ =1.24 (Tables 2, S1). This indicates that the relationships of osteocyte lacuna volume with genome size and ploidy level varies depending on phylogenetic affinities. Taxa that experienced a recent polyploidisation event (shown with arrowheads in Fig. 1, and in black on Fig. 10) belong to four different lineages: the sturgeon *Acipenser gueldenstaedtii*, the catostomid sucker *Catostomus commersoni*, the carp *Cyprinus carpio*, and several salmonid species (Fig. 1). We find that recent polyploid taxa tend to have the largest osteocyte lacunae and some of the largest genome sizes (Fig. 10). The carp *C. carpio* is an exception, with its moderately large osteocyte lacunae of the ribs and a relatively small genome compared to other polyploid species (note that it has relatively small osteocyte lacunae in the dentary; Fig. 10). Overall, much of the variation in genome size among our sample is driven by the occurrence of recent polyploidisation events.

Teleosts underwent an ancient genome duplication prior to their most recent common ancestor (Fig. 1), and non-teleost actinopterygian taxa (in white on Fig. 10) that did not experience this duplication event (to the exception of the sturgeon *A. gueldenstaedtii* that underwent a polyploidisation event specific to sturgeons), do not cluster together in our regression analyses (Fig. 10). Two of these taxa, the bichirs *Polypterus delhezi* and *Erpetoichthys calabaricus* have some of the largest genome sizes of all non-polyploid taxa, and have relatively large osteocyte lacunae. Compared to these, the gars *Atractosteus tropicus* and *Lepisosteus oculatus* have small genomes and smaller osteocyte lacunae. The bowfin *Amia calva* has a relatively small genome and large osteocyte lacunae compared to these taxa.

## Discussion

### Intra-bone variation

Our observations on variation within individual bones are consistent with the hypothesis that osteocyte lacuna volumes are related to bone patterns and therefore to growth rate. We show, for the first time both in 3D and in actinopterygians, that osteocyte lacunae vary in volume as much as they vary in shape and orientation among bone deposition patterns (Figs. 3–6). This is consistent with observations in other literature based on fossil and extant tetrapods (Baud, 1976; Marotti, 1979, 1981; de Ricqlès *et al*., 1991; Kerschnitzki *et al*., 2011; Cadena & Schweitzer, 2012; Sanchez *et al*., 2014, 2016; Stein & Prondvai, 2014; van Oers *et al*., 2015). Notably, osteocytes in the woven bone of the actinopterygians studied here are very large. Osteocytes are much smaller within areas of organised bone (i.e. parallel-fibered or lamellar bone).

In tetrapods, variation in osteocyte shapes and orientation results from changes in the extracellular collagen-fibre mesh (Marotti, 1979; Hernandez *et al*., 2004; Kerschnitzki *et al*., 2011; van Oers *et al*., 2015; Warshaw *et al*., 2017). This can be the consequence of different bone growth rates (Padian *et al*., 2001; Warshaw *et al*., 2017). In this case, the volumes of osteocytes greatly depends on those of their precursor osteoblasts (Marotti, 1976; Zambonin Zallone, 1977). Since osteoblasts secrete the osteoid that subsequently becomes bone, a faster rate of bone deposition is associated with larger osteoblasts (Marotti, 1976; de Ricqlès *et al*., 1991). For instance, woven bone tends to be associated with larger osteocytes and is deposited faster by larger osteoblasts (Baud, 1976; de Ricqlès *et al*., 1991; Remaggi *et al*., 1998; Cadena & Schweitzer, 2012; Stein & Prondvai, 2014; Grunmeier & D’Emic, 2019). Our observations on actinopterygian woven and organised bone tissues, therefore suggest that bone development in this group is regulated similarly to tetrapod bone as far the sizes of osteoblasts, and subsequently osteocytes, is concerned (see above; Fig. 4).

We also find that osteocyte lacuna volumes vary cyclically during deposition of periosteal bone (Fig. 5). Cell spaces are especially small in the regions preceding lines of arrested growth (e.g. carp rib, Fig. 5B; arowana rib, Fig. 5D), in the areas where visible collagen lamellae clearly indicate lamellar bone (Fig. 2D), and towards the outer surface of the bone in most specimens (e.g. Fig. 5A). This pattern probably corresponds to the cyclical bone growth described in sarcopterygians. The bones of stem- and crown-group tetrapods display growth marks that comprise “zones” as the result of an active metabolic period, and “annuli” as evidence of a slow-down in bone-growth, classically followed by lines of arrested growth (LAGs) which punctuate a lethargic period. LAGs are often visible in SRμCT images (e.g. Sanchez *et al*., 2016; Kamska *et al*., 2018). Cell-volume variation is associated with this cyclical pattern in sarcopterygians (Castanet *et al*., 1993; Sanchez *et al*., 2014). As zones, annuli and LAGs could be clearly recognised in some specimens of actinopterygians studied here (Fig. 5B, E) and because they correlate with cell volume fluctuations (Fig. 5A, D), we strongly suggest that cells in these actinopterygians undergo a cyclical fluctuation paralleling cyclical growth patterns. The smaller cell volumes observed in the periphery of the actinopterygian bones studied here coincides with decreasing growth rates during the acquisition of skeletal maturity, paralleling patterns seen in sarcopterygians.

Areas identified as secondary bone – from the presence of cementing lines of reversal and from conspicuous local changes in osteocyte orientation – exhibit notable differences in osteocyte lacuna volumes compared to the adjacent primary bone in some cases only (Fig. 6B). Differences occur only when the bone deposition pattern (i.e. organised or woven) is different. In these cases, osteocytes are almost always smaller in secondary than in the adjacent primary bone (Fig. 6D). This is consistent with observations of the Haversian mammalian bone, in which remodelling results in organised slowly-deposited and mechanically stronger secondary bone (such as lamellar bone) (Currey, 2002; van Oers *et al*., 2015). Secondary bone of the actinopterygians sampled here secondary bone may have deposited slowly for similar functional reasons related to mechanical strengthening.

### Intra-skeletal variation

Different bones of the same organism have different osteocyte lacuna volumes, even within a single bone pattern (e.g. within areas of organised bone). We find substantial differences in osteocyte lacuna volumes between the different skeletal elements of *Cyprinus carpio* (approximately three-fold variation; Fig. 7). In contrast, different regions within the same skeletal element do not show large differences in osteocyte volumes (Fig. 8). Despite the clarity of this pattern of variation among carp bones, there is no obvious explanation at present (Fig. 7). Bones that ossify early in ontogeny (e.g. vertebrae, operculum) do not seem to have larger osteocytes than those that ossify later in the development. Likewise, there is no obvious pattern in terms of bone size, position (cranial vs. postcranial bones), embryological origin (neural-crest derived vs. mesoderm) or type of ossification (dermal vs. endochondral). This finding is reminiscent of patterns in the emu *Dromaius novaehollandiae* (D’Emic & Benson, 2013), which shows a five-fold difference in lacuna volumes between bones of the skeleton. As for birds, the intra-skeletal variation in osteocyte volume observed in actinopterygians (Fig. 9) is probably explained by other factors.

### Inter-specific variation

We find greater than 9-fold variation in median osteocyte lacuna volumes among species, from 84.73 μm^3^ in the wolf herring *Chirocentrus dorab* to 784.95 μm^3^ in the salmonid *Stenodus leucichthys* for lacunae of parallel-fibered bone of the dentary. Species-specific growth rate could theoretically provide a good explanation for this inter-specific variation. Our analyses index growth rate using body length. However, this variable is not found to correlate with osteocyte lacuna volumes in our comparative analyses (Fig. 10). Instead, genome size has the highest explanatory power for lacuna volume amongst all biological parameters that we tested (Table 2). Even so, the explanatory power of the relationship is relatively low (R^2^ = 0.17 to 0.22 for dentary osteocyte lacunae; R^2^ = 0.43 to 0.49 for rib osteocyte lacunae), indicating considerable as-yet unexplained interspecific variation in osteocyte lacuna volumes.

Cell size covariation with genome size is a well-known phenomenon across the tree of life (Cavalier-Smith, 1982; Olmo, 1983; Gregory, 2000, 2001a; b, 2005; Mueller, 2015), for cells as diverse as erythrocytes (Olmo, 1983; Gregory, 2000, 2005; Hardie & Hebert, 2003), neurons (Roth *et al*., 1994), the stomatal guard cells of angiosperms (e.g. Beaulieu *et al*., 2008), and vertebrate osteocytes (predominantly in tetrapods; Organ *et al*., 2007, 2011; Organ & Shedlock, 2009; Montanari *et al*., 2011; D’Emic & Benson, 2013). Grunmeier & D’Emic (2019) recently demonstrated that this relationship holds for both static and dynamic osteogenesis in birds. In actinopterygians, the scaling between cell size and genome size is well-known in erythrocytes (Lay & Baldwin, 1999; Gregory, 2001b; Hardie & Hebert, 2003) but it is found here for the first time in actinopterygian osteocytes.

Another result of our study is that the increase in osteocyte volume is particularly evident in ray-finned fish taxa that experienced a recent polyploidisation event (Fig. 10). In addition to the teleost-specific whole genome duplication, several actinopterygian lineages experienced polyploidisation events (Leggatt & Iwama, 2003; Hardie & Hebert, 2004; Mable *et al*., 2011): non-teleosts such as sturgeons (Blacklidge & Bidwell, 1993; Birstein *et al*., 1997; Ludwig *et al*., 2001) and teleosts like catostomid suckers (Uyeno & Smith, 1972), the common carp *Cyprinus carpio* (David *et al*., 2003) and salmonids (Berthelot *et al*., 2014; Macqueen & Johnston, 2014). These polyploidisation events are thought to be relatively recent, for example Late Cretaceous (between 88 and 100 million years ago) in the case of salmonids (Macqueen & Johnston, 2014). Studies have suggested that increases in cell volumes may be more prominent for recent polyploidisation events due to the increase in cellular activity (e.g. transcription) that is necessary to comply with the doubling of genomic elements (Marshall *et al*., 2012). However, it is potentially less evident in older polyploidisation events because of genome-size reduction via loss of paralogous gene copies (Mueller, 2015). This seems to be the case with the teleost-specific genome duplication, an event which age is estimated to range between 316 and 226 million years ago (Santini *et al*., 2009; Inoue *et al*., 2015), and must have occurred by the first fossil appearance of crown-group teleosts in the Late Jurassic, approximately 150 million years ago (Hurley *et al*., 2007). Because of evolutionary genome size reduction since then (Brunet *et al*., 2006; Inoue *et al*., 2015), many teleosts in fact have smaller genomes than non-teleosts that did not experience this whole-genome duplication, such as polypterids (Fig. 10). These findings suggest that the fossil record of osteocytes might provide unique, and as-yet untapped, data on the history of genome evolution in actinopterygians.

## Conclusion

We provide the first three-dimensional overview of variation in osteocyte lacuna volumes within individuals and among species of ray-finned fishes (actinopterygians). This group comprises half of all vertebrate species but the study of their bone growth and microstructure has been relatively neglected. Our findings confirm patterns previously described from thin sections or (occasionally) three-dimensional data in other vertebrates:

1. Osteocyte lacunae vary significantly in volume within a given bone, and this variation most likely reflects changes in the rate of bone deposition.
2. This variation is mostly explained by three factors: bone deposition patterns (woven vs. organised bone tissues), bone remodelling and cyclical growth, and can thus be mitigated in comparative (inter-specific) studies by controlling the area of the bone that is sampled.
3. The volumes of osteocyte lacunae vary considerably from one bone to another across the skeleton (approximately three-fold variation among the bones of a carp). These results mandate that inter-specific studies aiming for meaningful comparison of osteocyte lacuna volumes should select homologous bones across the sampled taxa, and measure osteocytes from a consistent pattern of bone deposition (D’Emic & Benson, 2013; Grunmeier & D’Emic, 2019).

This study is one of the few to explore comparative three-dimensional variation in osteocyte lacunae volumes from SRμCT data, and as far as we know the first to adopt a multi-scale (e.g. from the intra-bone to the interspecific) approach in ray-finned fishes. SRμCT data have proven very advantageous for this endeavour: their high resolution and capacity to generate three-dimensional data considerably reduce uncertainties on shape and volume estimations, and the high throughput scanning setups and semi-automated measurement methods that we employed increase the amount of available data to a level way above that of comparable studies.

Our inter-specific comparisons corroborate studies on other taxa showing genome size to be the best predictor for the volume of osteocyte lacunae (e.g. D’Emic & Benson, 2013). However, the large variation in these volumes that we observe likely incorporates other biological signals outside of genome size, possibly including growth rates and metabolic rates (that we were not able to test directly). Furthermore, the relationship between osteocyte size and genome size varies among species according to phylogeny, though this variation is limited. Nevertheless, fossilised osteocyte lacuna volume may be especially useful to describe and detect large-scale changes in genome size through evolutionary history. These include high-magnitude jumps in genome size due to polyploidisation or ancient genome duplication events, or longterm trends in genome size increase or decrease without any change in ploidy level, as known in lungfishes (Thomson, 1972) or in urodeles (Laurin *et al*., 2016; Liedtke *et al*., 2018).

Osteocytes remain the best available proxy to study vertebrate genomic events in extinct taxa: they have been used to track genome size increase in lungfishes (Thomson, 1972) and its decrease in pterosaurs (Organ & Shedlock, 2009) and dinosaurs (Organ *et al*., 2007, 2009), as well as to estimate the ancestral genome size of tetrapods (Thomson & Muraszko, 1978; Organ *et al*., 2016), lissamphibians (Laurin *et al*., 2016) and amniotes (Organ *et al*., 2011). However, none of these studies has made use of X-ray μCT and resulting 3D datasets, which potentially introduces biases in measuring osteocyte lacuna volumes. Moreover, none so far have focused on ray-finned fishes, in which osteocytes preserved in ancient fossil representatives could be used to detect both the teleost-specific genome duplication event and their subsequent reduction in genome size, opening unprecedented avenues of research on their genome evolution in the deep time.

## Acknowledgements

The authors would like to thanks the curators and colleagues that provided access to the specimens we used: Matt Friedman and Doug Nelson (University of Michigan), Olga Otero (Université de Poitiers), Eileen Westwig (OUMNH) and in particular Philippe Béarez (MNHN) for allowing us to use his personal research collection. Chris Organ (Montana State University) provided comparative data on tetrapod osteocyte lacunae and genome sizes. T. Ryan Gregory (University of Guelph) is thanked for helping us to solve issues with access to the Animal Genome Size Database. Synchrotron beamtime was obtained via two proposals accepted at the ESRF (LS2614 and LS2758). Paul Tafforeau (ESRF) provided invaluable help and scientific input during the experiments. This work was supported by a grant from the Leverhulme Trust (RPG-2016-168), and by a Junior Research Fellowship at Wolfson College, University of Oxford.

