## Supplementary Information for "Three-dimensional characterisation of osteocyte volumes at multiple scales, and its relationship with bone biology and genome evolution in ray-finned fishes"

### 1. SRµCT scan setups

All specimens were characterised at the ID19 beamline of the ESRF (European Synchrotron Radiation Facility, Grenoble, France) using propagation phase contrast X-ray synchrotron micro computed tomography (PPC-SRµCT). We used four different setups to acquire data with a voxel size of 0.7 µm, changing the energy to adjust the dose intake on the sample and avoid movement during the scans.

*Detected energy of 19 keV*

Pink beam from an undulator (U17.6, gap 23 mm, 4 Al focussing lenses); indirect detector comprising a 10 µm GGG scintillator, a 10x Olympus microscope lens (Olympus Corporation, Tokyo, Japan) and PCO.edge 5.5; sample-detector distance was set to 20 mm. A 100 µm thick graphite rotating disk was placed before the sample, decreasing the coherence of the beam, effectively removing contribution from impurities on the Beryllium window. Tomographic acquisitions consisted of 2999 projection with an exposure time of 30 ms.

*Detected energy of 35 keV*

Pink beam from an undulator (U17.6, gap 23 mm, 4 Be focussing lenses); indirect detector (10 µm GGG scintillator, 10x Olympus microscope lens,PCO.edge 5.5); sample-detector distance set to 50 mm. Tomographic acquisitions consisted of 2999 projection with an exposure time of 60 ms.

*Detected energy of 105 keV*

Filtered white beam from a wiggler (gap 40 mm, 4.5 mm of copper, 70 Be focussing lenses); indirect detector (25 µm LuAG scintillator (Lutetium Aluminium Garnet), 10x Olympus microscope lens and PCO.edge Gold 4.2 sCMOS with USB3 camera link); Sample-detector distance was set to 200 mm. Tomographic acquisitions consisted of 2999 projection with an exposure time of 200 ms.

*Detected energy of 112 keV*

Filtered white beam from a wiggler (gap 38 mm, Cu 4 mm, 60 Be lenses); indirect detector (25 µm LuAG scintillator, 10x Olympus microscope lens,PCO.edge 4.2 sCMOS with USB3 camera link); sample-detector distance was set to 220 mm. Tomographic acquisitions consisted of 2999 projection with an exposure time of 200 ms.

**2. Supplementary figures and tables**

***Figure S1:*** Phylogenetic generalised least squares regression (pGLS) of median osteocyte lacuna volumes against genome sizes, common body length, maximum body weight and mature body length (all $\log_{10}$-transformed), using dentaries and the “max” threshold. Colour code as in Fig. 10.

***Figure S2:*** Phylogenetic generalised least squares regression (pGLS) of median osteocyte lacuna volumes against genome sizes, common body length, maximum body weight and mature body length (all $\log_{10}$-transformed), using dentaries and the “min” threshold. Colour code as in Fig. 10.

***Figure S3:*** Phylogenetic generalised least squares regression (pGLS) of median osteocyte lacuna volumes against genome sizes, common body length, maximum body weight and mature body length (all $\log_{10}$-transformed), using ribs and the “max” threshold. Colour code as in Fig. 10.

***Figure S4:*** Phylogenetic generalised least squares regression (pGLS) of median osteocyte lacuna volumes against genome sizes, common body length, maximum body weight and mature body length (all $\log_{10}$-transformed), using ribs and the “min” threshold. Colour code as in Fig. 10.

***Table S2:*** Detailed results of the phylogenetic generalised least squares regression (pGLS). 100 iterations have been performed, and the results are presented for the 25, 50 and 75% quantiles. Separate analyses were ran for dentaries and ribs, and for the “max” and “min” thresholds.
