## Supplementary figures and images for "Three-dimensional characterisation of osteocyte volumes at multiple scales, and its relationship with bone biology and genome evolution in ray-finned fishes"

### Figure S1

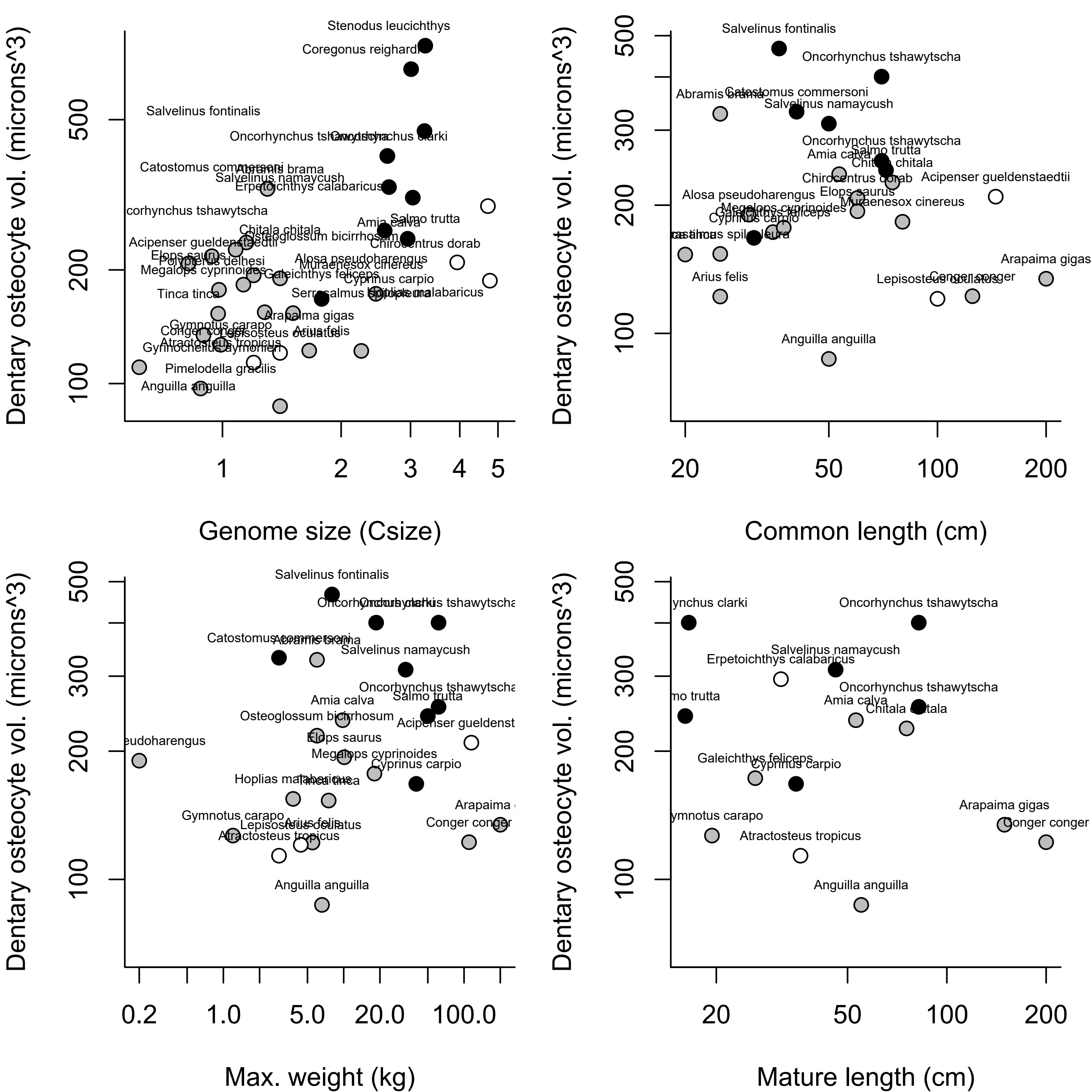

### Figure S2

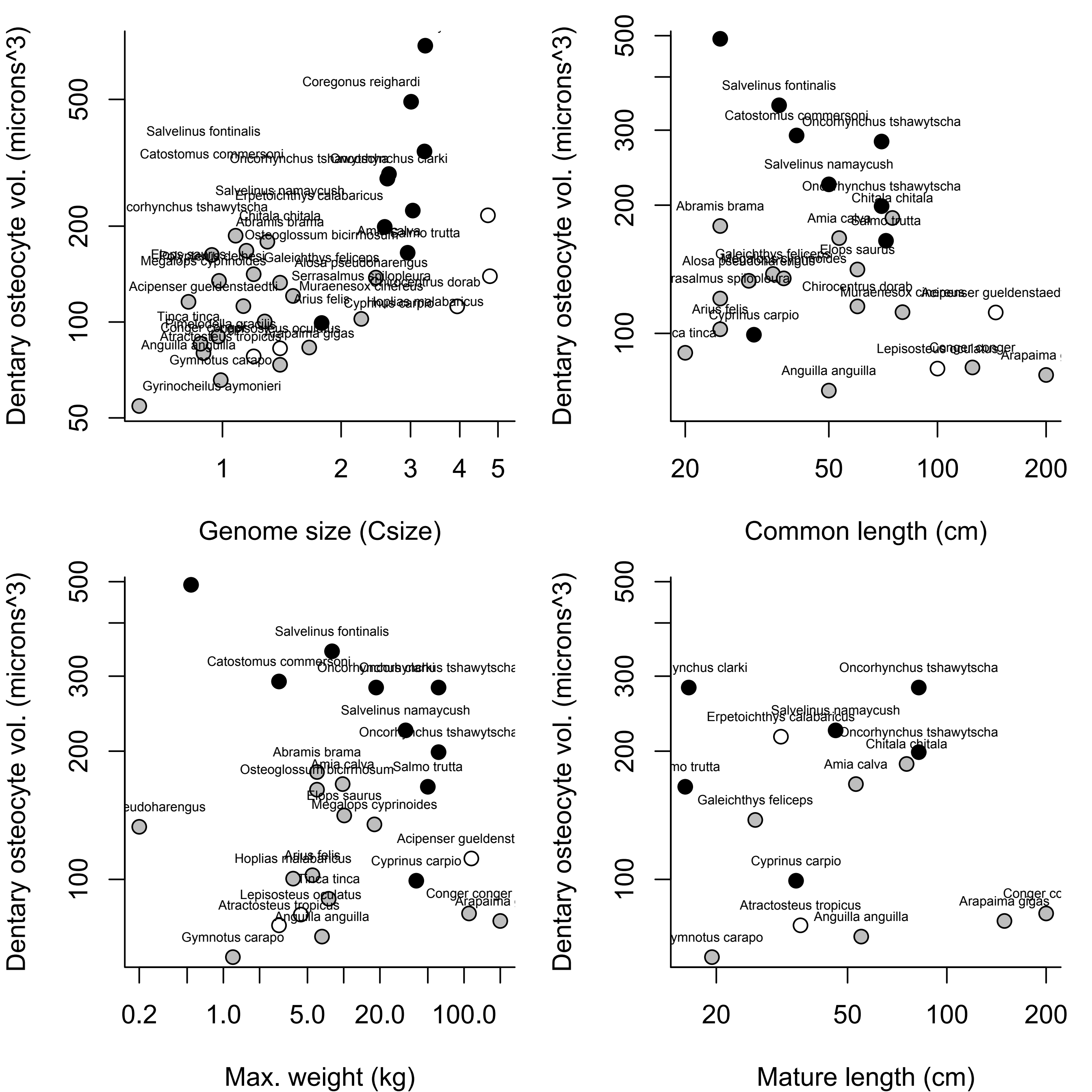

### Figure S3

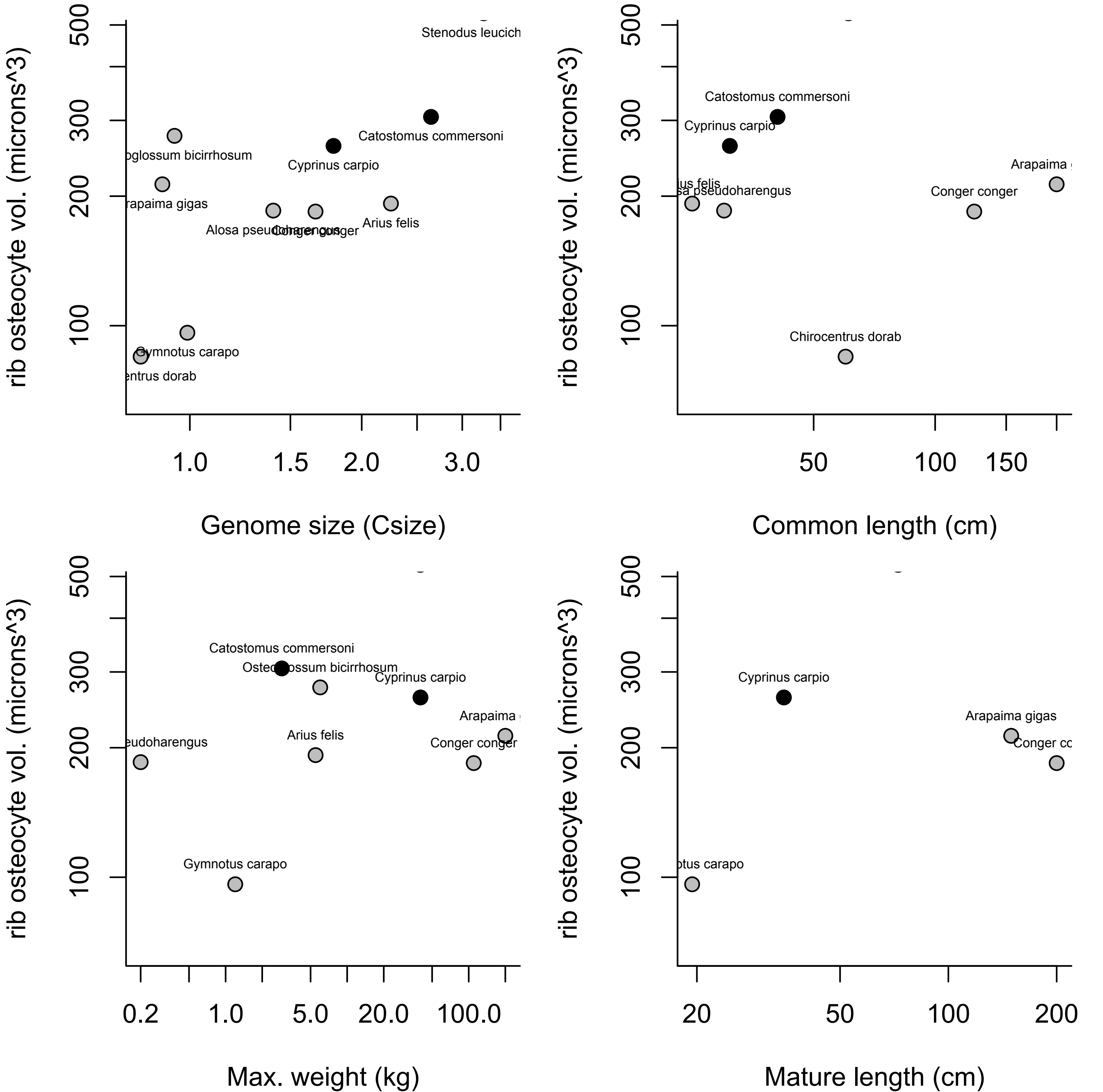

### Figure S4

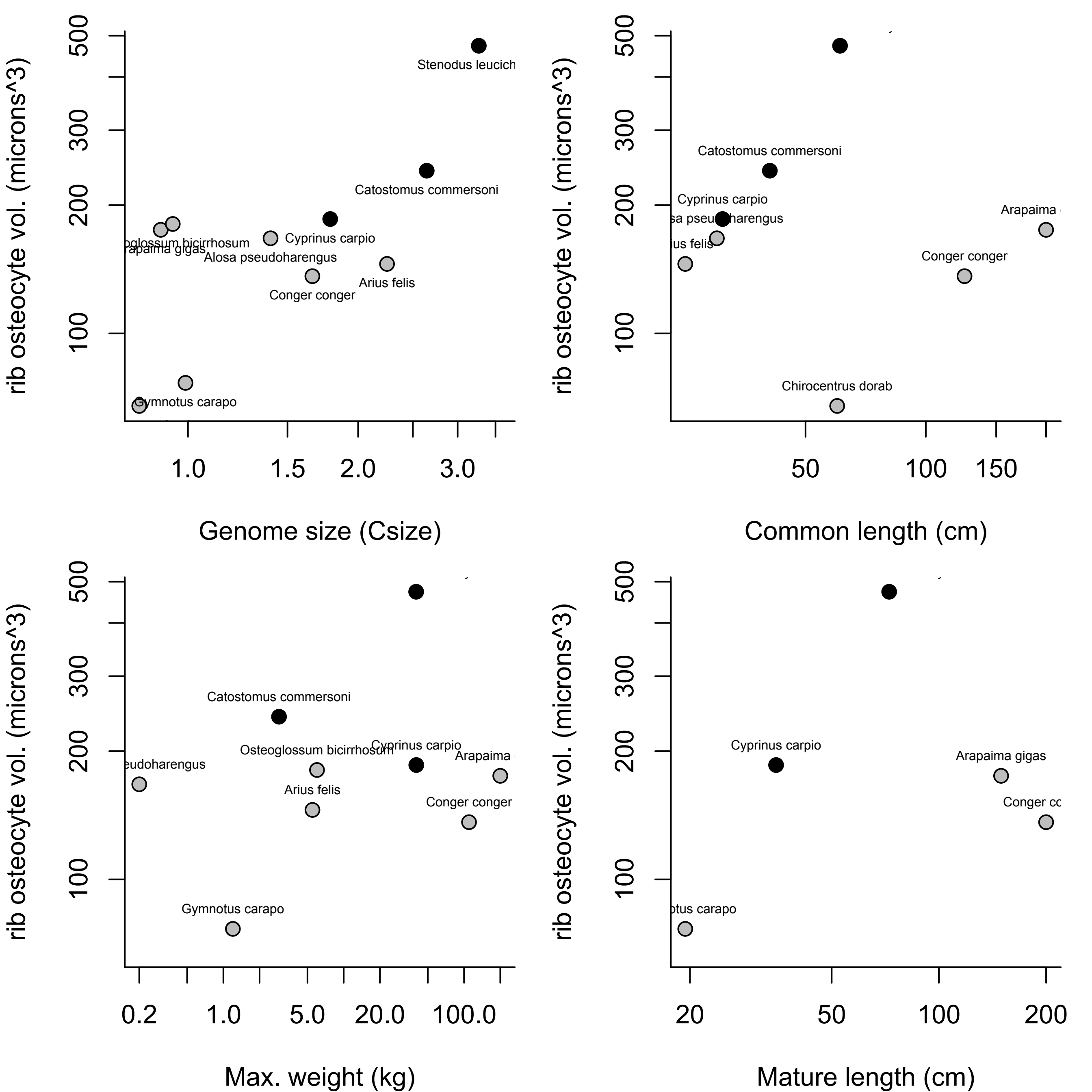
