## Supplementary material for "Three-dimensional characterisation of osteocyte volumes at multiple scales, and its relationship with bone biology and genome evolution in ray-finned fishes": Table S1

**Dentary, "max" threshold, 25% quantile**

| | model | AICc | $R^2$ | λ | Value | Std.Error | p-value |
|---|---|---|---|---|---|---|---|
| (Intercept) | osteocyte lacuna volume ~ genome size (C-value) | -9.593 | 0.146 | 0.393 | 2.163 | 0.074 | 0 |
| log10(Csize.temp) | | | | | 0.421 | 0.147 | 0.005 |
| (Intercept) | osteocyte lacuna volume ~ ploidy level | -9.033 | 0.14 | 0.054 | 2.215 | 0.039 | 0 |
| duplication.temp | | | | | 0.259 | 0.072 | 0.001 |
| (Intercept) | osteocyte lacuna volume ~ genome size + ploidy level | -6.649 | 0.141 | 0.012 | 2.184 | 0.04 | 0 |
| log10(Csize.temp) | | | | | 0.22 | 0.157 | 0.123 |
| duplication.temp | | | | | 0.176 | 0.088 | 0.046 |
| (Intercept) | osteocyte lacuna volume ~ genome size + body length | -4.871 | 0.08 | 0.425 | 2.066 | 0.223 | 0 |
| log10(length.temp) | | | | | -0.069 | 0.106 | 0.53 |
| log10(Csize.temp) | | | | | 0.414 | 0.151 | 0.006 |
| (Intercept) | osteocyte lacuna volume ~ body length + ploidy level | -4.687 | 0.068 | 0.108 | 2.261 | 0.197 | 0 |
| log10(length.temp) | | | | | -0.103 | 0.1 | 0.306 |
| duplication.temp | | | | | 0.266 | 0.076 | 0.002 |
| (Intercept) | osteocyte lacuna volume ~ body length | -2.282 | -0.078 | 0.492 | 2.11 | 0.248 | 0 |
| log10(length.temp) | | | | | -0.05 | 0.118 | 0.502 |
| (Intercept) | osteocyte lacuna volume ~ genome size + ploidy level + body length | -1.926 | 0.07 | 0.151 | 2.174 | 0.203 | 0 |
| log10(length.temp) | | | | | -0.088 | 0.1 | 0.4 |
| log10(Csize.temp) | | | | | 0.218 | 0.172 | 0.139 |
| duplication.temp | | | | | 0.167 | 0.096 | 0.069 |

**Dentary, "min" threshold, 25% quantile**

| | model | AICc | $R^2$ | λ | Value | Std.Error | p-value |
|---|---|---|---|---|---|---|---|
| (Intercept) | osteocyte lacuna volume ~ genome size (C-value) | -6.063 | 0.226 | 0.361 | 1.967 | 0.078 | 0 |
| log10(Csize.temp) | | | | | 0.53 | 0.159 | 0.002 |
| (Intercept) | osteocyte lacuna volume ~ genome size + body length | -2.99 | 0.176 | 0.404 | 2.035 | 0.234 | 0 |
| log10(length.temp) | | | | | -0.167 | 0.111 | 0.152 |
| log10(Csize.temp) | | | | | 0.555 | 0.162 | 0.002 |
| (Intercept) | osteocyte lacuna volume ~ ploidy level | -2.687 | 0.134 | 0.123 | 2.055 | 0.052 | 0 |
| duplication.temp | | | | | 0.26 | 0.084 | 0.003 |
| (Intercept) | osteocyte lacuna volume ~ genome size + ploidy level | -2.212 | 0.192 | 0.147 | 1.992 | 0.062 | 0 |
| log10(Csize.temp) | | | | | 0.36 | 0.186 | 0.048 |
| duplication.temp | | | | | 0.123 | 0.103 | 0.184 |
| (Intercept) | osteocyte lacuna volume ~ body length + ploidy level | 0.077 | 0.09 | 0.193 | 2.243 | 0.225 | 0 |
| log10(length.temp) | | | | | -0.186 | 0.111 | 0.108 |
| duplication.temp | | | | | 0.279 | 0.086 | 0.003 |
| (Intercept) | osteocyte lacuna volume ~ genome size + ploidy level + body length | 0.876 | 0.148 | 0.25 | 2.114 | 0.234 | 0 |
| log10(length.temp) | | | | | -0.177 | 0.111 | 0.131 |
| log10(Csize.temp) | | | | | 0.37 | 0.204 | 0.06 |
| duplication.temp | | | | | 0.137 | 0.114 | 0.194 |
| (Intercept) | osteocyte lacuna volume ~ body length | 3.002 | -0.067 | 0.439 | 2.104 | 0.267 | 0 |
| log10(length.temp) | | | | | -0.128 | 0.129 | 0.322 |

**Dentary, "max" threshold, 50% quantile (Table 2)**

| | model | AICc | $R^2$ | λ | Value | Std.Error | p-value |
|---|---|---|---|---|---|---|---|
| (Intercept) | osteocyte lacuna volume ~ genome size (C-value) | -8.888 | 0.17 | 0.438 | 2.167 | 0.079 | 0 |
| log10(Csize.temp) | | | | | 0.428 | 0.15 | 0.007 |
| (Intercept) | osteocyte lacuna volume ~ ploidy level | -8.147 | 0.144 | 0.094 | 2.22 | 0.044 | 0 |
| duplication.temp | | | | | 0.269 | 0.074 | 0.001 |
| (Intercept) | osteocyte lacuna volume ~ genome size + ploidy level | -6.267 | 0.153 | 0.097 | 2.186 | 0.048 | 0 |
| log10(Csize.temp) | | | | | 0.239 | 0.165 | 0.145 |
| duplication.temp | | | | | 0.182 | 0.092 | 0.053 |
| (Intercept) | osteocyte lacuna volume ~ genome size + body length | -4.174 | 0.098 | 0.484 | 2.289 | 0.227 | 0 |
| log10(length.temp) | | | | | -0.063 | 0.108 | 0.554 |
| log10(Csize.temp) | | | | | 0.433 | 0.153 | 0.008 |
| (Intercept) | osteocyte lacuna volume ~ body length + ploidy level | -4.074 | 0.085 | 0.17 | 2.418 | 0.202 | 0 |
| log10(length.temp) | | | | | -0.101 | 0.101 | 0.329 |
| duplication.temp | | | | | 0.267 | 0.077 | 0.002 |
| (Intercept) | osteocyte lacuna volume ~ body length | -1.502 | -0.074 | 0.525 | 2.38 | 0.255 | 0 |
| log10(length.temp) | | | | | -0.049 | 0.121 | 0.605 |
| (Intercept) | osteocyte lacuna volume ~ genome size + ploidy level + body length | -1.034 | 0.086 | 0.19 | 2.35 | 0.215 | 0 |
| log10(length.temp) | | | | | -0.083 | 0.103 | 0.411 |
| log10(Csize.temp) | | | | | 0.228 | 0.18 | 0.196 |
| duplication.temp | | | | | 0.182 | 0.101 | 0.07 |

**Dentary, "min" threshold, 50% quantile**

| | model | AICc | $R^2$ | λ | Value | Std.Error | p-value |
|---|---|---|---|---|---|---|---|
| (Intercept) | osteocyte lacuna volume ~ genome size (C-value) | -5.2 | 0.236 | 0.406 | 1.974 | 0.082 | 0 |
| log10(Csize.temp) | | | | | 0.547 | 0.164 | 0.002 |
| (Intercept) | osteocyte lacuna volume ~ genome size + body length | -2.043 | 0.198 | 0.54 | 2.248 | 0.242 | 0 |
| log10(length.temp) | | | | | -0.145 | 0.114 | 0.196 |
| log10(Csize.temp) | | | | | 0.572 | 0.166 | 0.002 |
| (Intercept) | osteocyte lacuna volume ~ ploidy level | -1.689 | 0.141 | 0.158 | 2.056 | 0.055 | 0 |
| duplication.temp | | | | | 0.274 | 0.087 | 0.004 |
| (Intercept) | osteocyte lacuna volume ~ genome size + ploidy level | -1.498 | 0.201 | 0.218 | 1.998 | 0.064 | 0 |
| log10(Csize.temp) | | | | | 0.38 | 0.198 | 0.054 |
| duplication.temp | | | | | 0.142 | 0.11 | 0.203 |
| (Intercept) | osteocyte lacuna volume ~ body length + ploidy level | 0.827 | 0.113 | 0.263 | 2.398 | 0.232 | 0 |
| log10(length.temp) | | | | | -0.173 | 0.115 | 0.131 |
| duplication.temp | | | | | 0.293 | 0.089 | 0.003 |
| (Intercept) | osteocyte lacuna volume ~ genome size + ploidy level + body length | 1.479 | 0.167 | 0.404 | 2.301 | 0.241 | 0 |
| log10(length.temp) | | | | | -0.159 | 0.114 | 0.162 |
| log10(Csize.temp) | | | | | 0.386 | 0.207 | 0.068 |
| duplication.temp | | | | | 0.146 | 0.117 | 0.235 |
| (Intercept) | osteocyte lacuna volume ~ body length | 4.196 | -0.052 | 0.473 | 2.338 | 0.279 | 0 |
| log10(length.temp) | | | | | -0.112 | 0.134 | 0.39 |

**Dentary, "max" threshold, 75% quantile**

| | model | AICc | $R^2$ | λ | Value | Std.Error | p-value |
|---|---|---|---|---|---|---|---|
| (Intercept) | osteocyte lacuna volume ~ genome size (C-value) | -8.215 | 0.184 | 0.476 | 2.168 | 0.081 | 0 |
| log10(Csize.temp) | | | | | 0.448 | 0.153 | 0.011 |
| (Intercept) | osteocyte lacuna volume ~ ploidy level | -7.244 | 0.147 | 0.15 | 2.221 | 0.048 | 0 |
| duplication.temp | | | | | 0.272 | 0.076 | 0.002 |
| (Intercept) | osteocyte lacuna volume ~ genome size + ploidy level | -5.146 | 0.164 | 0.168 | 2.187 | 0.056 | 0 |
| log10(Csize.temp) | | | | | 0.255 | 0.172 | 0.196 |
| duplication.temp | | | | | 0.19 | 0.095 | 0.061 |
| (Intercept) | osteocyte lacuna volume ~ genome size + body length | -3.494 | 0.12 | 0.506 | 2.301 | 0.234 | 0 |
| log10(length.temp) | | | | | 0.049 | 0.114 | 0.687 |
| log10(Csize.temp) | | | | | 0.446 | 0.158 | 0.014 |
| (Intercept) | osteocyte lacuna volume ~ body length + ploidy level | -2.186 | 0.09 | 0.184 | 2.42 | 0.214 | 0 |
| log10(length.temp) | | | | | -0.02 | 0.108 | 0.852 |
| duplication.temp | | | | | 0.275 | 0.08 | 0.002 |
| (Intercept) | osteocyte lacuna volume ~ body length | -0.824 | -0.062 | 0.567 | 2.385 | 0.27 | 0 |
| log10(length.temp) | | | | | 0.088 | 0.129 | 0.683 |
| (Intercept) | osteocyte lacuna volume ~ genome size + ploidy level + body length | 0.229 | 0.097 | 0.263 | 2.358 | 0.223 | 0 |
| log10(length.temp) | | | | | 0.004 | 0.109 | 0.954 |
| log10(Csize.temp) | | | | | 0.259 | 0.182 | 0.244 |
| duplication.temp | | | | | 0.185 | 0.104 | 0.108 |

**Dentary, "min" threshold, 75% quantile**

| | model | AICc | $R^2$ | λ | Value | Std.Error | p-value |
|---|---|---|---|---|---|---|---|
| (Intercept) | osteocyte lacuna volume ~ genome size (C-value) | -1.602 | 0.243 | 0.45 | 1.977 | 0.086 | 0 |
| log10(Csize.temp) | | | | | 0.566 | 0.17 | 0.003 |
| (Intercept) | osteocyte lacuna volume ~ ploidy level | 2.115 | 0.146 | 0.211 | 2.059 | 0.06 | 0 |
| duplication.temp | | | | | 0.29 | 0.089 | 0.004 |
| (Intercept) | osteocyte lacuna volume ~ genome size + ploidy level | 2.244 | 0.206 | 0.286 | 2 | 0.075 | 0 |
| log10(Csize.temp) | | | | | 0.395 | 0.201 | 0.078 |
| duplication.temp | | | | | 0.146 | 0.111 | 0.244 |
| (Intercept) | osteocyte lacuna volume ~ genome size + body length | 3.114 | 0.214 | 0.588 | 2.281 | 0.267 | 0 |
| log10(length.temp) | | | | | -0.034 | 0.13 | 0.796 |
| log10(Csize.temp) | | | | | 0.589 | 0.175 | 0.003 |
| (Intercept) | osteocyte lacuna volume ~ body length + ploidy level | 6.264 | 0.136 | 0.305 | 2.422 | 0.26 | 0 |
| log10(length.temp) | | | | | -0.096 | 0.13 | 0.472 |
| duplication.temp | | | | | 0.302 | 0.096 | 0.004 |
| (Intercept) | osteocyte lacuna volume ~ genome size + ploidy level + body length | 6.946 | 0.185 | 0.454 | 2.329 | 0.266 | 0 |
| log10(length.temp) | | | | | -0.064 | 0.129 | 0.625 |
| log10(Csize.temp) | | | | | 0.403 | 0.213 | 0.094 |
| duplication.temp | | | | | 0.156 | 0.123 | 0.246 |
| (Intercept) | osteocyte lacuna volume ~ body length | 8.965 | -0.044 | 0.507 | 2.37 | 0.311 | 0 |
| log10(length.temp) | | | | | 0.011 | 0.15 | 0.945 |

**Rib, "max" threshold, 25% quantile**

| | model | AICc | $R^2$ | λ | Value | Std.Error | p-value |
|---|---|---|---|---|---|---|---|
| (Intercept) | osteocyte lacuna volume ~ genome size (C-value) | 1.693 | 0.613 | 1.24 | 2.154 | 0.094 | 0 |
| log10(Csize.temp) | | | | | 1.015 | 0.217 | 0.002 |
| (Intercept) | osteocyte lacuna volume ~ genome size + ploidy level | 7.328 | 0.557 | 1.175 | 2.148 | 0.093 | 0 |
| log10(Csize.temp) | | | | | 0.863 | 0.265 | 0.014 |
| duplication.temp | | | | | 0.126 | 0.139 | 0.393 |
| (Intercept) | osteocyte lacuna volume ~ ploidy level | 7.559 | 0.304 | 0.518 | 2.23 | 0.083 | 0 |
| duplication.temp | | | | | 0.354 | 0.129 | 0.025 |
| (Intercept) | osteocyte lacuna volume ~ genome size + body length | 7.984 | 0.527 | 1.268 | 2.299 | 0.322 | 0 |
| log10(length.temp) | | | | | -0.065 | 0.134 | 0.644 |
| log10(Csize.temp) | | | | | 0.983 | 0.244 | 0.005 |
| (Intercept) | osteocyte lacuna volume ~ body length | 11.987 | -0.084 | 1.038 | 2.672 | 0.532 | 0.001 |
| log10(length.temp) | | | | | -0.156 | 0.241 | 0.536 |
| (Intercept) | osteocyte lacuna volume ~ body length + ploidy level | 13.255 | 0.199 | 0.614 | 2.309 | 0.437 | 0.001 |
| log10(length.temp) | | | | | -0.036 | 0.203 | 0.864 |
| duplication.temp | | | | | 0.357 | 0.14 | 0.038 |
| (Intercept) | osteocyte lacuna volume ~ genome size + ploidy level + body length | 15.027 | 0.475 | 1.239 | 2.354 | 0.337 | 0 |
| log10(length.temp) | | | | | -0.094 | 0.143 | 0.536 |
| log10(Csize.temp) | | | | | 0.821 | 0.298 | 0.033 |
| duplication.temp | | | | | 0.144 | 0.153 | 0.383 |

**Rib, "min" threshold, 25% quantile**

| | model | AICc | $R^2$ | λ | Value | Std.Error | p-value |
|---|---|---|---|---|---|---|---|
| (Intercept) | osteocyte lacuna volume ~ genome size (C-value) | 1.859 | 0.495 | 1.21 | 2.032 | 0.093 | 0 |
| log10(Csize.temp) | | | | | 1.093 | 0.221 | 0.001 |
| (Intercept) | osteocyte lacuna volume ~ genome size + ploidy level | 7.685 | 0.411 | 1.142 | 2.03 | 0.093 | 0 |
| log10(Csize.temp) | | | | | 0.946 | 0.276 | 0.011 |
| duplication.temp | | | | | 0.11 | 0.142 | 0.463 |
| (Intercept) | osteocyte lacuna volume ~ ploidy level | 8.195 | 0.049 | 0.364 | 2.117 | 0.081 | 0 |
| duplication.temp | | | | | 0.356 | 0.131 | 0.026 |
| (Intercept) | osteocyte lacuna volume ~ genome size + body length | 8.267 | 0.376 | 1.181 | 2.049 | 0.338 | 0 |
| log10(length.temp) | | | | | -0.006 | 0.144 | 0.966 |
| log10(Csize.temp) | | | | | 1.075 | 0.251 | 0.004 |
| (Intercept) | osteocyte lacuna volume ~ body length | 10.027 | -0.142 | -0.601 | 1.947 | 0.33 | 0 |
| log10(length.temp) | | | | | 0.106 | 0.173 | 0.557 |
| (Intercept) | osteocyte lacuna volume ~ body length + ploidy level | 13.765 | -0.081 | 0.115 | 1.92 | 0.434 | 0.003 |
| log10(length.temp) | | | | | 0.091 | 0.206 | 0.671 |
| duplication.temp | | | | | 0.346 | 0.135 | 0.038 |
| (Intercept) | osteocyte lacuna volume ~ genome size + ploidy level + body length | 15.597 | 0.287 | 1.13 | 2.092 | 0.362 | 0.001 |
| log10(length.temp) | | | | | -0.027 | 0.157 | 0.868 |
| log10(Csize.temp) | | | | | 0.917 | 0.322 | 0.029 |
| duplication.temp | | | | | 0.118 | 0.157 | 0.48 |

**Rib, "max" threshold, 50% quantile**

| | model | AICc | $R^2$ | λ | Value | Std.Error | p-value |
|---|---|---|---|---|---|---|---|
| (Intercept) | osteocyte lacuna volume ~ genome size (C-value) | 1.693 | 0.613 | 1.24 | 2.154 | 0.094 | 0 |
| log10(Csize.temp) | | | | | 1.015 | 0.217 | 0.002 |
| (Intercept) | osteocyte lacuna volume ~ genome size + ploidy level | 7.328 | 0.557 | 1.175 | 2.148 | 0.093 | 0 |
| log10(Csize.temp) | | | | | 0.863 | 0.265 | 0.014 |
| duplication.temp | | | | | 0.126 | 0.139 | 0.393 |
| (Intercept) | osteocyte lacuna volume ~ ploidy level | 7.559 | 0.304 | 0.518 | 2.23 | 0.083 | 0 |
| duplication.temp | | | | | 0.354 | 0.129 | 0.025 |
| (Intercept) | osteocyte lacuna volume ~ genome size + body length | 7.984 | 0.527 | 1.268 | 2.299 | 0.322 | 0 |
| log10(length.temp) | | | | | -0.065 | 0.134 | 0.644 |
| log10(Csize.temp) | | | | | 0.983 | 0.244 | 0.005 |
| (Intercept) | osteocyte lacuna volume ~ body length | 11.987 | -0.084 | 1.038 | 2.672 | 0.532 | 0.001 |
| log10(length.temp) | | | | | -0.156 | 0.241 | 0.536 |
| (Intercept) | osteocyte lacuna volume ~ body length + ploidy level | 13.255 | 0.199 | 0.614 | 2.309 | 0.437 | 0.001 |
| log10(length.temp) | | | | | -0.036 | 0.203 | 0.864 |
| duplication.temp | | | | | 0.357 | 0.14 | 0.038 |
| (Intercept) | osteocyte lacuna volume ~ genome size + ploidy level + body length | 15.027 | 0.475 | 1.239 | 2.354 | 0.337 | 0 |
| log10(length.temp) | | | | | -0.094 | 0.143 | 0.536 |
| log10(Csize.temp) | | | | | 0.821 | 0.298 | 0.033 |
| duplication.temp | | | | | 0.144 | 0.153 | 0.383 |

**Rib, "min" threshold, 50% quantile**

| | model | AICc | $R^2$ | λ | Value | Std.Error | p-value |
|---|---|---|---|---|---|---|---|
| (Intercept) | osteocyte lacuna volume ~ genome size (C-value) | 1.859 | 0.495 | 1.21 | 2.032 | 0.093 | 0 |
| log10(Csize.temp) | | | | | 1.093 | 0.221 | 0.001 |
| (Intercept) | osteocyte lacuna volume ~ genome size + ploidy level | 7.685 | 0.411 | 1.142 | 2.03 | 0.093 | 0 |
| log10(Csize.temp) | | | | | 0.946 | 0.276 | 0.011 |
| duplication.temp | | | | | 0.11 | 0.142 | 0.463 |
| (Intercept) | osteocyte lacuna volume ~ ploidy level | 8.195 | 0.049 | 0.364 | 2.117 | 0.081 | 0 |
| duplication.temp | | | | | 0.356 | 0.131 | 0.026 |
| (Intercept) | osteocyte lacuna volume ~ genome size + body length | 8.267 | 0.376 | 1.181 | 2.049 | 0.338 | 0 |
| log10(length.temp) | | | | | -0.006 | 0.144 | 0.966 |
| log10(Csize.temp) | | | | | 1.075 | 0.251 | 0.004 |
| (Intercept) | osteocyte lacuna volume ~ body length | 10.027 | -0.142 | -0.601 | 1.947 | 0.33 | 0 |
| log10(length.temp) | | | | | 0.106 | 0.173 | 0.557 |
| (Intercept) | osteocyte lacuna volume ~ body length + ploidy level | 13.765 | -0.081 | 0.115 | 1.92 | 0.434 | 0.003 |
| log10(length.temp) | | | | | 0.091 | 0.206 | 0.671 |
| duplication.temp | | | | | 0.346 | 0.135 | 0.038 |
| (Intercept) | osteocyte lacuna volume ~ genome size + ploidy level + body length | 15.597 | 0.287 | 1.13 | 2.092 | 0.362 | 0.001 |
| log10(length.temp) | | | | | -0.027 | 0.157 | 0.868 |
| log10(Csize.temp) | | | | | 0.917 | 0.322 | 0.029 |
| duplication.temp | | | | | 0.118 | 0.157 | 0.48 |

**Rib, "max" threshold, 75% quantile**

| | model | AICc | $R^2$ | λ | Value | Std.Error | p-value |
|---|---|---|---|---|---|---|---|
| (Intercept) | osteocyte lacuna volume ~ genome size (C-value) | 1.693 | 0.613 | 1.24 | 2.154 | 0.094 | 0 |
| log10(Csize.temp) | | | | | 1.015 | 0.217 | 0.002 |
| (Intercept) | osteocyte lacuna volume ~ genome size + ploidy level | 7.328 | 0.557 | 1.175 | 2.148 | 0.093 | 0 |
| log10(Csize.temp) | | | | | 0.863 | 0.265 | 0.014 |
| duplication.temp | | | | | 0.126 | 0.139 | 0.393 |
| (Intercept) | osteocyte lacuna volume ~ ploidy level | 7.559 | 0.304 | 0.518 | 2.23 | 0.083 | 0 |
| duplication.temp | | | | | 0.354 | 0.129 | 0.025 |
| (Intercept) | osteocyte lacuna volume ~ genome size + body length | 7.984 | 0.527 | 1.268 | 2.299 | 0.322 | 0 |
| log10(length.temp) | | | | | -0.065 | 0.134 | 0.644 |
| log10(Csize.temp) | | | | | 0.983 | 0.244 | 0.005 |
| (Intercept) | osteocyte lacuna volume ~ body length | 11.987 | -0.084 | 1.038 | 2.672 | 0.532 | 0.001 |
| log10(length.temp) | | | | | -0.156 | 0.241 | 0.536 |
| (Intercept) | osteocyte lacuna volume ~ body length + ploidy level | 13.255 | 0.199 | 0.614 | 2.309 | 0.437 | 0.001 |
| log10(length.temp) | | | | | -0.036 | 0.203 | 0.864 |
| duplication.temp | | | | | 0.357 | 0.14 | 0.038 |
| (Intercept) | osteocyte lacuna volume ~ genome size + ploidy level + body length | 15.027 | 0.475 | 1.239 | 2.354 | 0.337 | 0 |
| log10(length.temp) | | | | | -0.094 | 0.143 | 0.536 |
| log10(Csize.temp) | | | | | 0.821 | 0.298 | 0.033 |
| duplication.temp | | | | | 0.144 | 0.153 | 0.383 |

**Rib, "min" threshold, 75% quantile**

| | model | AICc | $R^2$ | λ | Value | Std.Error | p-value |
|---|---|---|---|---|---|---|---|
| (Intercept) | osteocyte lacuna volume ~ genome size (C-value) | 1.859 | 0.495 | 1.21 | 2.032 | 0.093 | 0 |
| log10(Csize.temp) | | | | | 1.093 | 0.221 | 0.001 |
| (Intercept) | osteocyte lacuna volume ~ genome size + ploidy level | 7.685 | 0.411 | 1.142 | 2.03 | 0.093 | 0 |
| log10(Csize.temp) | | | | | 0.946 | 0.276 | 0.011 |
| duplication.temp | | | | | 0.11 | 0.142 | 0.463 |
| (Intercept) | osteocyte lacuna volume ~ ploidy level | 8.195 | 0.049 | 0.364 | 2.117 | 0.081 | 0 |
| duplication.temp | | | | | 0.356 | 0.131 | 0.026 |
| (Intercept) | osteocyte lacuna volume ~ genome size + body length | 8.267 | 0.376 | 1.181 | 2.049 | 0.338 | 0 |
| log10(length.temp) | | | | | -0.006 | 0.144 | 0.966 |
| log10(Csize.temp) | | | | | 1.075 | 0.251 | 0.004 |
| (Intercept) | osteocyte lacuna volume ~ body length | 10.027 | -0.142 | -0.601 | 1.947 | 0.33 | 0 |
| log10(length.temp) | | | | | 0.106 | 0.173 | 0.557 |
| (Intercept) | osteocyte lacuna volume ~ body length + ploidy level | 13.765 | -0.081 | 0.115 | 1.92 | 0.434 | 0.003 |
| log10(length.temp) | | | | | 0.091 | 0.206 | 0.671 |
| duplication.temp | | | | | 0.346 | 0.135 | 0.038 |
| (Intercept) | osteocyte lacuna volume ~ genome size + ploidy level + body length | 15.597 | 0.287 | 1.13 | 2.092 | 0.362 | 0.001 |
| log10(length.temp) | | | | | -0.027 | 0.157 | 0.868 |
| log10(Csize.temp) | | | | | 0.917 | 0.322 | 0.029 |
| duplication.temp | | | | | 0.118 | 0.157 | 0.48 |
